# Characterization of BvlzGluc, a Novel Antifungal β-Glucanase from *Bacillus velezensis*, with Potential Agricultural and Industrial Applications

**DOI:** 10.1101/2025.01.23.634490

**Authors:** D. Vela-Corcia, A.M. Colino-Palomino, A. de Vicente, A. Pérez García, D. Romero

## Abstract

β-glucanases play a critical role in degrading β-glucans, which are essential components of fungal cell walls that maintain fungal structure and physiology. This study characterizes BvlzGluc, a novel β-glucanase produced by *Bacillus velezensis* CECT 8237, which has potent antifungal properties. BvlzGluc specifically targets the β-glucan layer of *Botrytis cinerea*, causing significant structural malformations, impaired fungal development, and reduced virulence. These effects weaken the fungal cell wall, increasing its susceptibility to environmental stress and reducing its capacity to infect host plants. Notably, BvlzGluc is a noncanonical β-glucanase belonging to glycoside hydrolase family 16 (GH16) but presents unique structural features that distinguish it from typical members of this family, suggesting potential reclassification. Its physicochemical stability and simple structural design increase its suitability for scalable production, positioning it as a promising candidate for agricultural and industrial applications requiring efficient β-glucan degradation. This work builds on previous studies of microbial antifungal mechanisms, providing new insights into bacterial‒fungal interactions and reinforcing the potential of bacterial enzymes in sustainable pathogen management and advancing biotechnological innovations.

## 1. Introduction

β-glucans are the primary polysaccharides found in the fungal cell wall and play essential roles in providing structural integrity and rigidity. These polysaccharides consist of β-1,3 and β-1,6 linkages that form a fibrillar network, which contributes to the overall strength and resilience of the fungal cell wall. β-1,3-glucan forms a helical structure that acts as a scaffold, while the β-1,6 linkages connect these helices, creating a highly cross-linked matrix. This combination of linkages provides both flexibility and mechanical support, allowing for the fungal cell wall to resist environmental stresses and osmotic pressure [1,2]. β-glucans are critical in fungal physiology, particularly in maintaining the integrity of the cell wall during processes such as budding, hyphal growth, and spore formation. Their synthesis and remodeling are tightly regulated by fungal cells, with β-glucanases and other enzymes involved in modulating the polymerization and depolymerization of β-glucan chains, ensuring dynamic adaptation to various environmental conditions [1].

β-glucanases are a group of glycoside hydrolases (GHs) that catalyze the conversion of β-glycosidic bonds in β-glucans into smaller oligosaccharides, such as cello-oligosaccharides and, occasionally, glucose. These enzymes are classified into various types based on their specific activity, including endo-β-1,4-glucanase (EC 3.2.1.4), endo-β-1,3(4)-glucanase (EC 3.2.1.6), and endo-β-1,3-glucanase/laminarinase (EC 3.2.1.39), among others, as categorized by the CAZy database [3]. They receive great interest in industries dedicated to biofuel production, where lignocellulosic biomass must be degraded into simpler sugars (e.g., glucose and xylose) before being refined into biofuels. By breaking down these complex polymers, glucanases help unlock renewable energy from plant-derived materials, making them valuable for sustainable energy solutions [4–7]. Additionally, they have been associated with crop protection and plant defense against fungal pathogens since they hydrolyze β-glucans in fungal cell walls, thus playing a crucial role in weakening structural barriers and, ultimately, the integrity of the pathogen [8]. This degradation exposes chitin, another key component of the fungal cell wall, making it more recognizable to the plant immune system. Upon exposure, chitin triggers plant defense mechanisms, such as the production of antimicrobial compounds and the activation of systemic acquired resistance (SAR), which limits fungal spread and infection [9–12]. *Bacillus velezensis* has emerged as a significant player because of its potent antifungal activities. The presence of β-glucanases in *B. velezensis* and their ability to suppress fungal infections make these enzymes crucial components in plant protection strategies [13,14]. Diverse β-glucanases have been described in members of this bacterial species and characterized for their roles in antagonistic interactions with fungi. [15][16,17].

In this study, we identified and characterized a novel β-glucanase, BvlzGluc, produced by *Bacillus velezensis* CECT 8237. BvlzGluc effectively degrades the β-glucan layer of fungal cell walls, compromising structural integrity and causing severe malformations that inhibit fungal development and reduce disease incidence. Our structural and functional analyses revealed that BvlzGluc has unique characteristics distinct from those of other β-glucanases in the glycoside hydrolase family, suggesting that it represents a new type of β-glucanase. These findings underscore the importance of bacterial factors in antagonistic interactions between biocontrol agents and pathogens, furthering innovative, sustainable agricultural and biotechnological pathogen control strategies that extend beyond traditional bacterial applications.

## 2. Materials and methods

### 2.1. Plant material

Melon plants (*Cucumis melo*; cultivar Rochet Panal) were grown from seeds (Semillas Fitó) at 25 °C and 60% relative humidity under fluorescent and incandescent light at a photofluency rate of ∼ 120 μmol m^-2^ s^-1^ and a 12/12 h photoperiod.

### 2.2. Fungal strain—growth and inoculation of plant material

*B. cinerea* isolate B05.10 was cultured on potato dextrose agar (PDA, Oxoid) in a controlled-environment chamber at 20 °C under illumination with fluorescent light at a photo fluency rate of 12 μmol m^-2^ s^-1^ and a 12/12 h photoperiod. Conidia were harvested from the light-grown culture in sterile distilled water containing 0.001% (v/v) Triton X-100 (J.T. Baker) and filtered through a 40-μm cell strainer to remove the remaining hyphae. Cell biology studies were carried out on germinated conidia grown in potato dextrose broth (PDB, Scharlau) inoculated with a spore suspension and incubated at 28 °C for 24 h at 150 rpm. *B. cinerea* plant inoculation was performed as previously described [18]. Briefly, the conidial suspension was adjusted to 10^5^ conidia mL^-1^ in half-strength filtered (0.45 μm) grape juice (100% pure organic). Melon plants that were 5–6 weeks old were used, and each leaf was inoculated with 5-μl droplets of conidial suspension (5·10^2^ conidia). The plants were covered with a plastic dome and placed in the growth chamber. At 72 h postinoculation, the leaves were imaged, and the lesion size was analyzed using ImageJ 2.0 software.

### 2.3. Stress analysis of B. cinerea B05.10: identifying active fractions and critical concentration estimation

A stress analysis of *B. cinerea* B05.10 was performed to identify the clone supernatants with antimicrobial activity. To do so, all the clones were grown in LB medium at 37 °C for 24 h at 150 rpm. Afterward, the supernatants were collected, filter sterilized and subsequently incubated with *B. cinerea* germinated spores in a 96-well microtiter plate. Then, reactive oxygen species (ROS) production was analyzed using 10 mM hydrogen peroxide as a positive control. ROS detection was performed with dihydrorhodamine 123 (DHR123, Sigma‒Aldrich). The fluorescence was recorded using a plate reader (FluoStar Omega, BMG LabTech) with excitation filters at 485 nm and emission at 520 nm, and readings were taken every 10 minutes over a 24-hour period. The experiment was stopped when the positive control became saturated.

### 2.4. Sequence analysis

A search was conducted in the fosmid cloned fragments for proteins that could be secreted. For this, a previously described workflow was followed [19]. Briefly, proteins with a signal peptide were searched using SignalP (https://services.healthtech.dtu.dk/service.php?SignalP-5.0), those without transmembrane domains were identified using TMHMM (https://services.healthtech.dtu.dk/service.php?TMHMM-2.0), and proteins predicted to have extracellular subcellular localization were selected using DeepLoc (https://services.healthtech.dtu.dk/service.php?DeepLoc-1.0). Finally, a comprehensive search for conserved protein domains was conducted on the selected proteins to identify functional motifs that may be critical for their activity (https://toolkit.tuebingen.mpg.de/tools/hhpred).

Multiple sequence alignment was conducted via the CLC Main Workbench (Aarhus). Phylogenetic analysis was performed by MEGA software (v.11, Kumar Lab, Temple University, Philadelphia, USA) with MUSCLE as the alignment algorithm. Phylogenetic tree analysis was subsequently carried out using the maximum likelihood statistical method, which is based on the Poisson correction model. The test to assess the phylogeny used was performed by the bootstrap method with 1000 bootstrap replications.

### 2.5. In vitro expression and purification of B. velezensis CECT 8237 BvlzGluc

The primers 7685F44-F (5’-AAAAAGCAGGCTCTATGCGTAAAAAAATCTTAATGGCACTC-3’) and 7685F44-R (5’-AGAAAGCTGGGTTTATCGATTAATTTTAAAAGAGTTTGTCG-3’) were used to amplify BAMY6639_07685 (GenBank acc. no. AMQ71412) coding sequence, the amplified fragment was cloned and inserted into the entry vector pDONR207 and then transferred to the destination vector pDEST17, which incorporates a six His-tag at the N-terminus of the proteins, using the Gateway^®^ cloning technology (Invitrogen).

For BvlzGluc expression, the plasmid pDEST17 containing the coding sequence was transformed into chemically competent *E. coli* BL21-AI cells (Invitrogen) using heat shock. Protein expression was induced with the addition of 10 mM arabinose (Sigma‒ Aldrich) at an OD_600 nm_ of 0.4. The cells were incubated at 37 °C for 3 h and harvested by centrifugation. The cell pellet was frozen in liquid nitrogen and stored at −80 °C overnight to increase the yield of protein recovery.

For protein purification, the cell pellet was thawed, resuspended in lysis buffer (1x CellLytic^TM^ B-Cell Lysis Reagent [Sigma‒Aldrich], 1 mM protease inhibitor phenylmethylsulfonyl fluoride (PMSF) and 0.2 mg mL^-1^ Lysozyme) and incubated at room temperature for 1 h; the remaining cells were disrupted by sonication on ice using a UP100H sonicator (Hielscher). The crude lysate was clarified by centrifugation at 8000 × g for 10 min.

The resulting supernatant was passed through a 0.45-μm filter prior to protein purification via affinity chromatography using an AKTA Start FPLC system (GE Healthcare). The lysate was loaded into a HisTrap HP 5 mL column (GE Healthcare) previously equilibrated with binding buffer (50 mM Na_3_PO_4_ [pH 8], 0.5 M NaCl, and 10 mM imidazole). The protein was eluted with elution buffer (50 mM Na_3_PO_4_ [pH 8], 0.5 M NaCl, 500 mM imidazole). Next, the purified protein was loaded into a HiPrep 26/10 desalting column (GE Healthcare), and the buffer was exchanged for 20 mM Tris-HCl, pH 8, and 50 mM NaCl. Finally, the protein was concentrated by ultrafiltration using Pierce^®^ Concentrators 20 K MWCO (Thermo Scientific) and stored at −20 °C until analysis. Prior to further analysis, the expressed proteins were characterized by western blotting. For immunoblot analysis, purified protein was electrophoresed on 12% SDS‒PAGE gels and electrotransferred onto polyvinylidene difluoride (PVDF) membranes using the Trans-Blot Turbo electrophoretic transfer cells (Bio‒Rad). The blots were probed with a 1:1000 dilution of a rabbit monoclonal anti-His-tag antibody (Rockland). The membranes were then incubated with a 1:20,000 dilution of a horseradish peroxidase-conjugated anti-rabbit antibody (Bio-Rad), and the bands were visualized by chemiluminescent detection via an enhanced chemiluminescence (ECL) western blotting analysis system (Thermo Scientific).

### 2.6. Germination rate and germ tube length measurement

To estimate the germination rate and measure germ tube length, *B. cinerea* spores were first prepared by collecting a spore suspension as described above, followed by adjusting the spore concentration. The spores were plated on water agar media in sterile Petri dishes in the absence or presence of BvlzGluc and spread evenly before incubation at 25 °C. Germination was monitored at regular intervals using a microscope, and a spore was considered germinated when the germ tube exceeded the spore diameter. The germination rate was determined by counting the total number of spores and the number of spores that germinated in randomly selected fields of view. Data on the germination rate and germ tube length were recorded and averaged, with experiments conducted in triplicate to ensure reproducibility.

### 2.7. Protein modeling and molecular docking

AlphaFold3 [20] was used for automated protein tertiary structure modeling of the *B. velezensis* CECT 8237 BvlzGluc protein. To identify potential binding sites of β-glucan (PubChem ID: 439262) to the BvlzGluc protein, automated molecular docking and thermodynamic analysis were performed using the web-based SwissDock program [www.swissdock.ch/docking] [21]. SwissDock predicts the possible molecular interactions between a target protein and small molecule on the basis of the docking algorithm EADock DSS [22]. The docking was performed using the “Accurate” parameter at otherwise default parameters, with no region of interest defined (blind docking). Binding energies were estimated by using CHARMM (Chemistry at HARvard Macromolecular Mechanics), a molecular simulation program implemented within SwissDock software, and the most favorable energies were evaluated by Fast Analytical Continuum Treatment of Solvation (FACTS). Finally, the energy results were scored and ranked by *full fitness* (kcal mol^−1^), and spontaneous binding was exhibited by the estimated Gibbs free energy ΔG (kcal mol^−1^). The negative values of ΔG support the assertion that the binding process is highly spontaneous. Modeling and docking results were visualized using UCSF Chimera v1.8 software.

### 2.8. Transmission and scanning electron microscopy

*B. cinerea* was fixed in 2.5% (v/v) glutaraldehyde and 4% (v/v) paraformaldehyde in 0.1 M phosphate buffer (PBS) overnight at 4 °C. The samples were postfixed in 1% osmium tetroxide solution in PBS for 90 min at room temperature, followed by washing with PBS and 15 min of stepwise dehydration in an ethanol series (30%, 50%, 70%, 90% and 100% twice). Between the 50% and 70% steps, the samples were incubated in a 2% uranyl acetate solution in 50% ethanol at 4 °C overnight. After dehydration, the samples were gradually embedded in low-viscosity Spurr’s resin (resin:ethanol, 1:1, 4 h; resin:ethanol, 3:1, 4 h; and pure resin overnight). The sample blocks were embedded in capsule molds containing pure resin for 72 h at 70 °C. The samples were imaged under an FEI Tecnai G2 20 TWIN transmission electron microscope at an accelerating voltage of 80 kV. The images were acquired using TIA FEI Imaging Software v.4.14.

For scanning electron microscopy, after ethanol dehydration, the samples were dried with a Bal-Tec CPD 030 critical point dryer. The dried samples were coated with a thin layer of gold using a Leica EM SCD050 coater before viewing under a JEOL JSM-6490 LV microscope.

### 2.9. Chitin and chitosan visualization

To evaluate the effects of BvlzGluc on the fungal cell wall, both chitin and chitosan were visualized. For chitin visualization, *Botrytis* cells that were untreated or incubated with 0.5 μM BvlzGluc for 24 hours were stained with 0.05 mg mL^-1^ WGA-Alexa Fluor 488^®^ conjugate in PBS for 1 hour at room temperature. After staining, the cells were washed three times with PBS containing Tween 20 before visualization.

For chitosan visualization, Eosin Y was employed, as it binds selectively to chitosan without showing affinity for chitin [23]. Following incubation with 0.5 μM BvlzGluc, the cells were washed twice with 1 mL of McIlvaine’s buffer (0.2 M Na₂HPO₄ and 0.1 M citric acid, pH 6.0), resuspended in 500 μL of the same buffer, and stained with 30 μL of Eosin Y (5 mg mL⁻¹ stock; Sigma) at room temperature in the dark for 10 minutes, followed by two washes with 1 mL of McIlvaine’s buffer to remove excess dye. Finally, the cells were resuspended in 500 μL of McIlvaine’s buffer.

The samples were imaged using a Zeiss LSM880 confocal microscope equipped with a Plan-apochromatic 63x/1.4 oil immersion objective. Chitin and chitosan were visualized with excitation at 488 nm, and images were acquired for further analysis.

### 2.10. Fungal cell wall extraction

Germinated spores of *B. cinerea* were captured by filtration through a 40-µm filter (Corning) to collect all the biomass, which was subsequently resuspended in 1 mL of lysis buffer (10 mM Tris-HCl pH 7.4, 1 mM PMSF). A 3-mm tungsten carbide bead (Qiagen) was added. The material was then disrupted in a TissueLyser II (Qiagen) at maximum speed with a frequency of 30 cycles s^-1^ for 30 minutes. This step was conducted to disrupt the cell wall. After this step, the sample was centrifuged at 1000 × g for 10 minutes at 4 °C, the supernatant was discarded, and the pellet was washed with 1 mL of solution A (1 mM PMSF). The mixture was subsequently centrifuged again at 1000 × g for 10 minutes at 4 °C, after which the pellet was washed with 1 mL of solution B (5% NaCl, 1 mM PMSF). This step was repeated, changing to solution C (2% NaCl, 1 mM PMSF) and 1 mL of solution D (1% NaCl, 1 mM PMSF). Finally, the pellet was weighed and resuspended in 100 µL of solution A. In a 96-well plate, the cell wall purification (100 µg mL^-1^) was combined with AMQ71412.1 protein at a concentration of 20 µg mL^-1^. The mixture was incubated at 28 °C with agitation for 24 hours. Afterward, the results were analyzed via MALDI‒TOF mass spectrometry.

### 2.11. Carbohydrate sedimentation assay

The carbohydrate sedimentation assay was performed as previously described [24]. Briefly, 0.5 µM BvlzGluc in 20 mM Tris (pH 8.0) was mixed with 1.5 mg of chitin (NEB), chitosan (Sigma‒Aldrich) or β-glucan (Sigma‒Aldrich) suspensions and incubated at room temperature for 2 h on an orbital shaker at 350 rpm. The same amount of protein in Tris buffer without added carbohydrates was used as a negative control. After centrifugation (5 min, 13,000 g), the supernatant was collected, and the pellet was washed three times with 800 µl of 20 mM Tris (pH 8.0) prior to resuspension in 2x Laemmli buffer with the addition of 5% β-mercaptoethanol as a reducing agent (Invitrogen). The presence of BvlzGluc in different fractions was determined by western blot analysis, as described above.

### 2.12. Immunofluorescence microscopy

*B. cinerea* cells incubated with 0.5 μM BvlzGluc for 24 h were fixed with 100% acetone for 10 min at -20 °C. After being washed with 1× PBS, the cells were blocked with blocking solution (3% (w/v) BSA and 0.2% (v/v) Triton X-100) for 60 min. Finally, the cells were stained for immunofluorescence using a 1:50 rabbit monoclonal anti-β-glucan primary antibody (Thermo Fisher) followed by a 1:200 GFP-conjugated goat anti-rabbit secondary antibody (Thermo Fisher). Images were obtained via visualization of the samples using a Zeiss LSM880 confocal microscope with a Plan-apochromatic 63x/1.4 oil immersion objective and acquisition with excitation at 488 nm. The cells were counterstained with the lipophilic dye FM 4-64 (Thermo Fisher) to stain early endosomes.

### 2.13. Enzyme activity assay

The activity of β-glucanase was assessed using the 3,5-dinitrosalicylic acid (DNS) method [25]. For this procedure, 0.5 μM BvlzGluc was incubated with 60 μL of 0.5% (w/v) barley β-glucan at 50 °C and pH 5.0 for 10 minutes. Then, 80 μL of DNS reagent was added, and the mixture was boiled for 5 minutes. After cooling to room temperature, the absorbance was measured at 540 nm using a spectrophotometer. One unit (U) of β-glucanase activity was defined as the amount of enzyme required to release 1 μmol of glucose-equivalent reducing sugar per minute. The enzyme concentration was determined with a Qubit, and 0.5 μM purified enzyme was used for physicochemical characterization.

### 2.14. Enzyme characterization

To establish the optimal pH, purified BvlzGluc was incubated with 0.5% barley β-glucan in various pH buffers, such as citrate‒phosphate buffer (pH 3.0–8.0), Tris‒HCl buffer (pH 8.0–9.0), and glycine‒NaOH buffer (pH 9.0–10.0), at 50 °C for 10 minutes, after which the glucanase activity was assessed. To investigate the pH stability of BvlzGluc, the enzyme was preincubated in buffers with a pH range of 3.0 to -10.0 for 1 hour at 25 °C, and the residual activities were measured. The enzyme activities before preincubation were designated 100%. To determine the optimal temperature for BvlzGluc, the enzyme was incubated with 0.5% barley β-glucan in citrate-phosphate buffer (pH 5.0) for 10 minutes at temperatures ranging from 20 °C to 90 °C.

### 2.15. Statistical analysis

GraphPad Prism v.10.0.0 (GraphPad Software, Boston, Massachusetts, USA) was used for statistical analysis of the experimental data. When the data were normally distributed and the sample variances were equal, Student’s *t* tests were performed with Welch’s correction. In all other cases, the Mann‒Whitney Rank Sum test was performed. For multiple comparisons, one-way analysis of variance (ANOVA) was performed when the equal variance test was passed. In all other cases, one-way ANOVA on ranks was performed (Kruskal‒Wallis significant difference test). Significance was accepted at P < 0.05.

## 3. Results and discussion

### 3.1. BvlzGluc reduces Botrytis spore germination and induces severe hyphal malformation

To identify protein determinants involved in the fungicidal activity of *B. velezensis* CECT 8237, a genomic library was constructed into the fosmid pCC2FOS™. A total of 400 clones were isolated, achieving full genome coverage of *Bacillus* at a depth of 4x. Compared with those from untreated plants or those treated with H_2_O_2_, cell-free supernatants from 12 different clones of the genome library induced a significant ROS response in response to *B. cinerea* spore suspensions (Supplementary Figure 1). Positive clones were sequenced and analyzed to identify secreted extracellular proteins with potential fungicidal properties. This analysis considered the search for signal peptides, the absence of transmembrane domains, and extracellular localization. The obtained sequences were also compared with known databases to pinpoint specific genes or gene clusters responsible for antifungal activity (Supplementary Table 1). One of the fosmids contained a genomic fragment with the locus BAMY6639_07685 that met all the aforementioned analysis criteria. The deduced amino acid sequence (GenBank acc. no. WP_060674061) annotated as a hypothetical protein was selected as a candidate for further analysis. The sequence of this protein shares significant homology with that of glycoside hydrolases, which are enzymes known for their role in the hydrolysis of glycosidic bonds, which was a reason to tentatively designate it as BvlzGluc to reflect its putative glucosidase activity.

The biological effects of BvlzGluc on *Botrytis cinerea*, specifically on fungal spore germination, were studied. The protein was heterogeneously expressed in *E. coli* and purified to homogeneity before further experiments were performed. The protein strongly inhibited the germination of fungal spores that produced abnormally short, twisted germ tubes with minimal elongation (Fig. 1a). Eventually, some spores could germinate but still produced significantly fewer germ tubes (Fig. 1b). Untreated spores (Fig. 1a, inset) presented normal and extended germ tubes, indicating healthy fungal growth. In addition, the external application of the protein to mature hyphae caused severe morphological deformations consisting of significant hyphal shortening accompanied by an increase in branching and pronounced vacuolization (Fig. 1c). This structural disruption suggested that the protein interferes with the normal growth and organization of fungal hyphae, resulting in abnormal cellular processes and compromising the overall integrity of the fungal network [26–28]. These malformations suggest that the protein exerts a dual inhibitory effect, impairing both the initial stages of fungal development and the integrity of mature structures.

**Figure 1.**
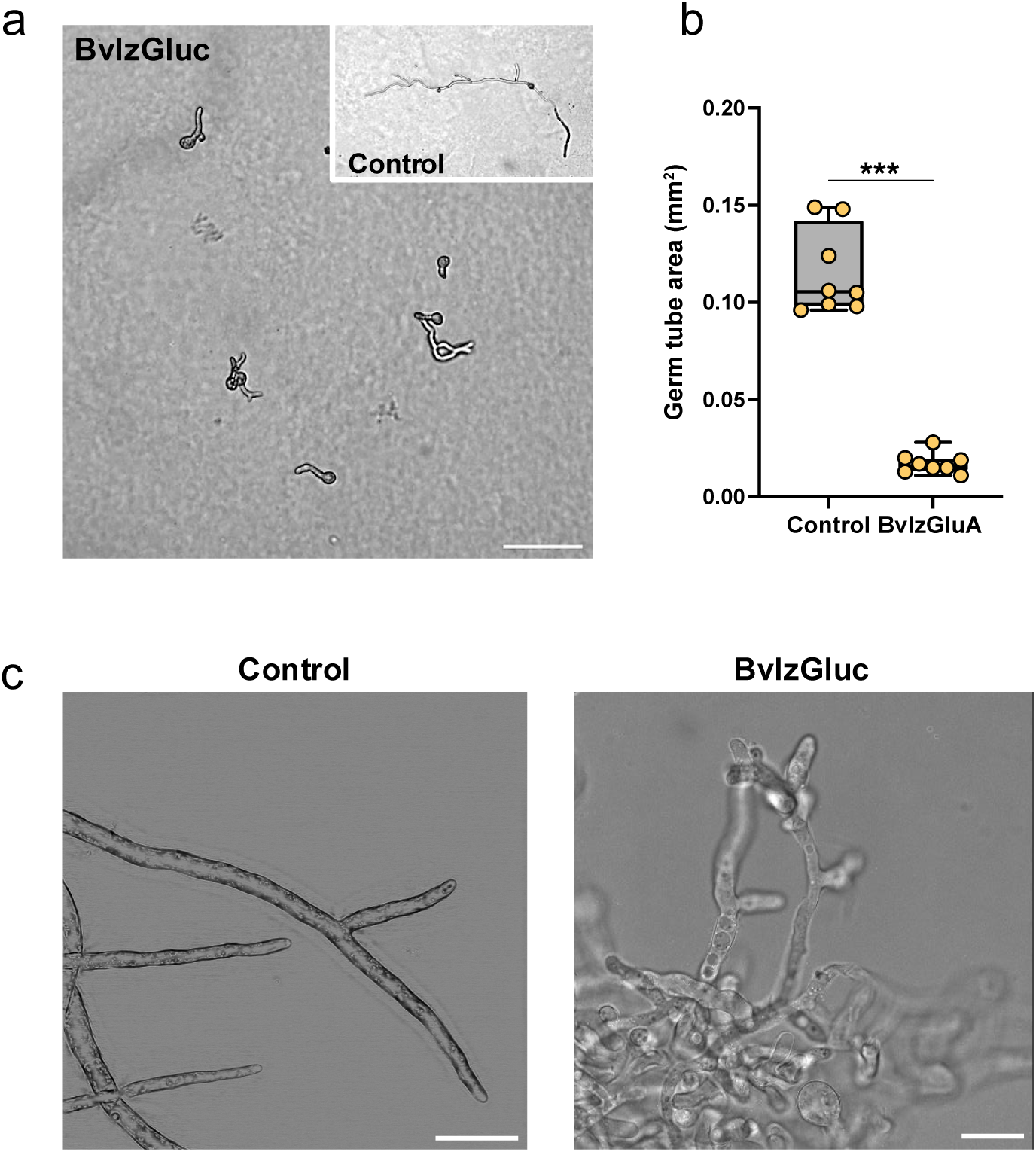
Impact of BvlzGluc on *Botrytis cinerea* development. **a**. The germination of *B. cinerea* spores was significantly inhibited upon exposure to BvlzGluc. In addition to the reduction in germination rates, the length of the germ tubes also considerably decreased. Scale bar, 20 μm. **b**. Whisker plot showing the quantification of the germ tube area. All measurements (yellow dots), medians (black line), and minimum and maximum values (whisker ends) are represented. The datasets did not pass the Shapiro‒Wilk test for normality (P > 0.05) and were compared using a nonparametric two-tailed Mann‒ Whitney test, with three asterisks indicating significant differences at P < 0.001. **c**. The effect of BvlzGluc on the development of *B. cinerea* was profound, leading to abnormal hyphal growth. The protein disrupted normal fungal development, resulting in the formation of short, highly branched hyphae. This aberrant growth pattern suggests that BvlzGluc interferes with the mechanisms that regulate hyphal elongation and branching. Scale bar, 30 μm.

The inhibition of fungal spore germination triggered by bacterial β-glucanase has been previously described in *B. subtilis* CW14 against *Aspergillus ochraceus* [29] and *B. velezensis* CE [17]. These findings suggest that the inhibitory activity of BvlzGluc may share mechanistic similarities with that of other β-glucanases in targeting fungal pathogens by disrupting spore development and germination processes, further highlighting its potential as an antifungal agent.

### 3.2. BvlzGluc disrupts the outer layer of the fungal cell wall

A hypothesis for the phenotypic malforming of hyphae is the disruptive effect of fungal cell surfaces [30–32]. Scanning electron microscopy analysis revealed the presence of an extracellular matrix surrounding the hyphae, which normally formed an interconnected network (Fig. 2A, top) that was clearly absent after treatment with BvlzGluc (Fig. 2a, bottom). This extracellular matrix is found in many fungi, both filamentous and nonfilamentous, and plays a crucial role in maintaining structural cohesion and communication between hyphae [33,34]. Transmission electron microscopy analysis of thin sections of fungal hyphae revealed that cell wall fragments were visibly released from the fungal structure (Fig. 2B, bottom, asterisk). No signs of disorganization or fragmentation of the fungal cell walls of the hyphae were observed after treatment with the buffer used for protein suspension (Fig. 2b, top). These anatomical deformities suggest that the protein might disrupt the outer layer of the cell wall and the extracellular matrix that holds the hyphae together. The loss of this matrix likely weakens the structural integrity of the fungal network, further contributing to the antifungal effect of BvlzGluc [35,36].

**Figure 2.**
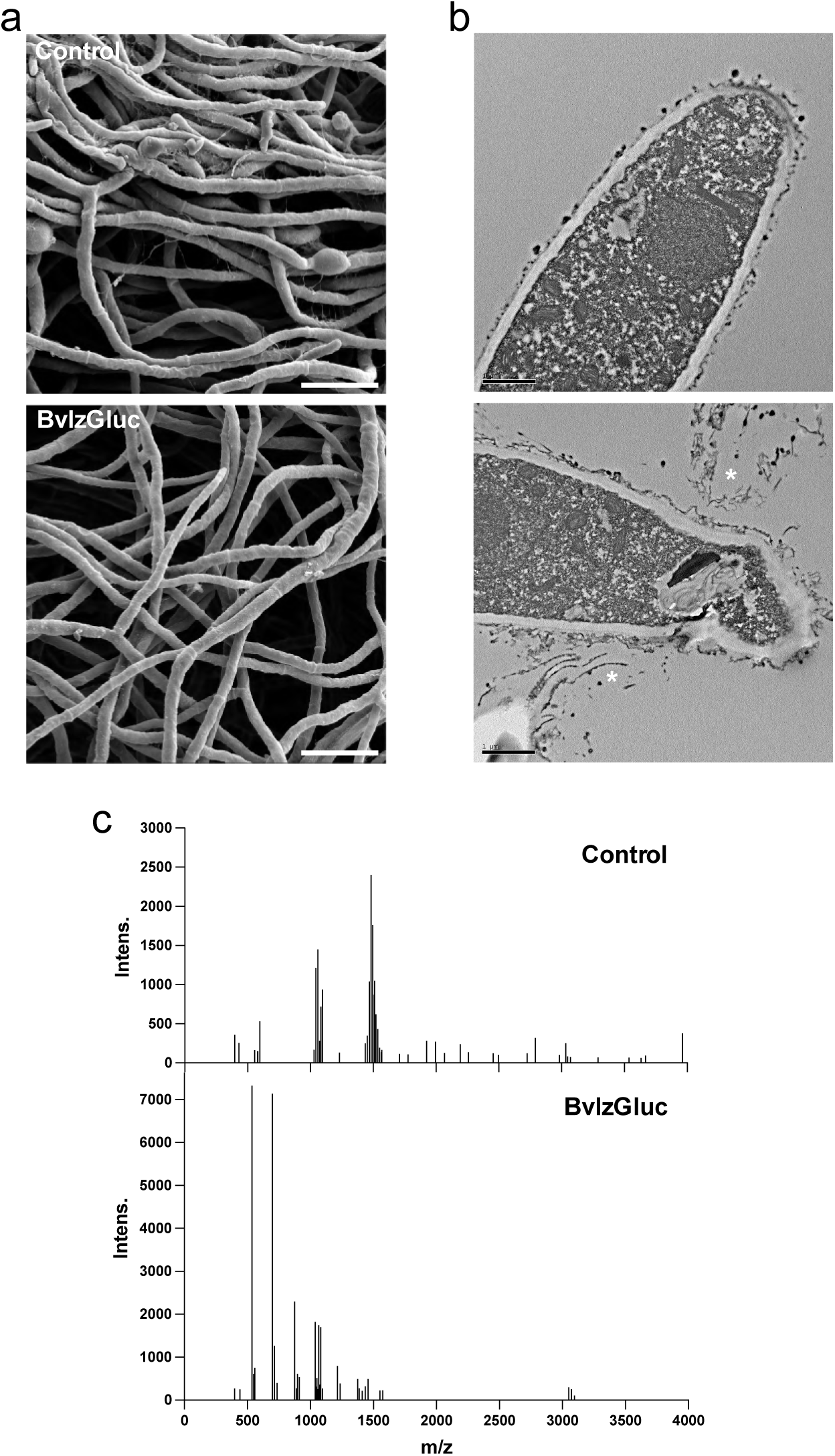
Impact of BvlzGluc on *Botrytis cinerea* hyphal integrity. **a**. Scanning electron micrographs showing the *B. cinerea* extracellular matrix. BvlzGluc manifests remarkable activity on the extracellular matrix fibers that normally connect fungal hyphae. Upon treatment with BvlzGluc, these connective fibers were absent and coexisted with disruption of the hyphal network. The absence of these extracellular matrix components suggests that BvlzGluc interferes with key structural proteins or polysaccharides, further highlighting its potential as a targeted antifungal agent. Scale bar, 20 μm. **b**. Transmission electron micrographs showing clear morphological alterations in samples treated with BvlzGluc compared with untreated *B. cinerea* hyphae. Upon treatment with BvlzGluc, notable changes in the fungal cell wall structure were observed, particularly the release of fragments from the outer layer of the hyphal cell wall (white asterisk). Scale bar, 1 μm. **c**. Mass spectrometry analysis of *B. cinerea* cell wall extracts from untreated and BvlzGluc-treated samples. In the untreated samples, high-molecular-weight peaks corresponding to the long chains of polysaccharides that form the fungal cell wall were detected. However, after treatment with BvlzGluc, these peaks completely disappeared, indicating a significant breakdown of the complex polysaccharide architecture of the cell wall. Instead, low-molecular-weight peaks, corresponding to oligomers and smaller fragments of degraded polysaccharides, emerged. This shift in molecular weight suggests that BvlzGluc catalyzes the hydrolysis of long polysaccharide chains into smaller and less complex derivatives.

To verify the specificity of the protein over certain components of the fungal cell wall and extracellular matrix, the purified cell wall was exposed to the protein and analyzed via mass spectrometry analysis. The chromatogram of untreated hyphae contained traces corresponding to high-molecular-weight polysaccharide chains, as expected for intact fungal cell walls. Treatment with the protein severely reduced the number of high-molecular-weight compounds and promoted the appearance of lower-molecular-weight compounds (Fig. 2c). These results suggest that the enzymatic degradation of cell wall polymers might be driven via the cleavage of glycosidic bonds and the transformation of macromolecules into oligomers [37,38], confirming the modification or dismantling of fungal cell wall structures.

### 3.3. BvlzGluc degrades fungal β-glucan, weakening cell wall integrity and reducing Botrytis cinerea virulence

The effect of BvlzGluc on *B. cinerea* was visualized *in vivo* via immunocytochemistry using a fungal anti-β-glucan-specific antibody. In the control sample, the β-glucan layer was clearly detected on the outermost region of the fungal cell wall. The fluorescence signal was continuous throughout the entire hypha, with higher intensity at the hyphal apex, indicating an intact and organized β-glucan layer structure. However, when the hyphae were treated with the protein BvlzGluc, the fluorescent signal associated with the β-glucan layer was no longer visible as a continuous structure. Fragments of the β-glucan layer adhered to the outer region of the cell wall, and additional β-glucan fragments were scattered randomly along the hyphae (Fig. 3a). This observation indicated that the protein had significant degrading activity on the β-glucan layer, disrupting the structural integrity of the cell wall and provoking a reduction in rigidity while gaining flexibility, which is vital for fungal growth and infection [29,39–41]. This degradation likely weakens the structural integrity of the cell wall, potentially reducing the ability of the fungus to grow and infect. To assess the biological impact of removing the β-glucan layer, the virulence of *Botrytis* was evaluated by measuring the lesion area on melon plant leaves. The lesion area in the presence of BvlzGluc was significantly smaller than that in the untreated control (Fig. 3b). On the basis of these findings, we reasoned that the degradation of the outer β-glucan layer may have a dual impact on reducing *Botrytis* virulence: i) directly destabilizing fungal integrity and viability and ii) indirectly triggering the immune response of the plant via exposure of the inner regions of the fungal cell wall [42–44].

**Figure 3.**
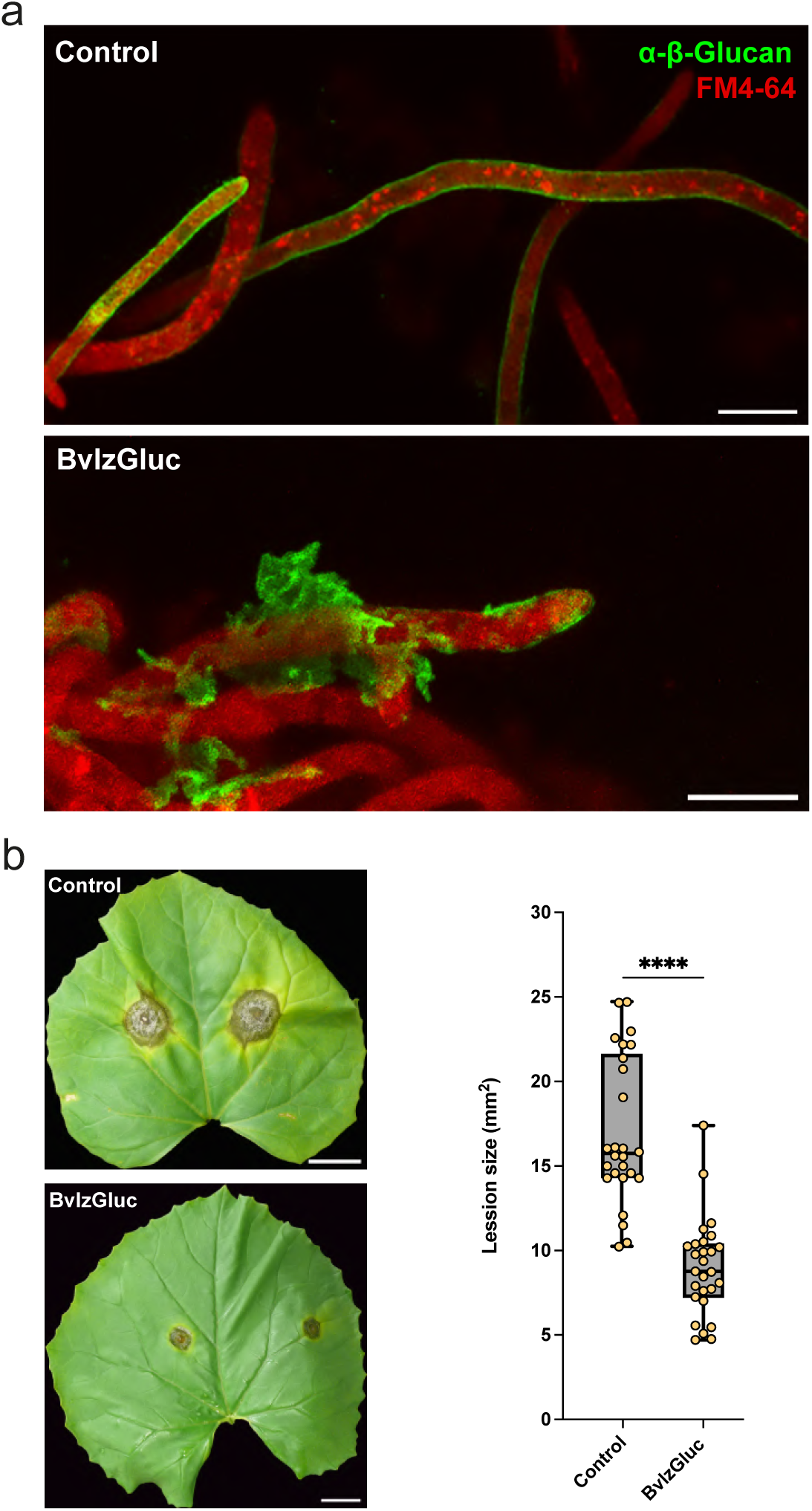
Effect of BvlzGluc on *Botrytis cinerea* infection in plants. **a**. Immunocytochemical analysis using an anti-fungal monoclonal primary antibody and a GFP-conjugated anti-rabbit secondary antibody demonstrated the normal distribution of β-glucans in *Botrytis cinerea* hyphae. In the control samples, the β-glucans formed a regular, continuous, and uniform layer within the cell wall, which is crucial for maintaining cell wall integrity and structural support in the fungus. However, upon treatment with BvlzGluc, significant disruptions in the β-glucan layer were observed. These discontinuities suggest that BvlzGluc interferes with the synthesis or maintenance of β-glucans, likely weakening the fungal cell wall and increasing its vulnerability to external stress. **b**. Lesion size triggered by *B. cinerea* on melon leaves after 72 hpi was reduced by 80% upon treatment with BvlzGluc. The whisker plot shows all the measurements (orange dots), medians (black line), and minimum and maximum values (whisker ends). The datasets did not pass the Shapiro‒Wilk test for normality (P > 0.05) and were compared using a nonparametric two-tailed Mann‒Whitney test, with quadruple asterisks indicating significant differences at P < 0.0001.

### 3.4. BvlzGluc specifically targets β-glucan through a noncanonical mechanism

To specifically determine the target of this protein, separate suspensions of each cell wall component (β-glucans, chitin, or chitosan) were prepared and incubated with the protein. After the incubation period, the samples were centrifuged to separate the pellet (containing nondegraded or insoluble material) from the supernatant (containing soluble or degraded components). Western blot analysis using anti-His tag antibodies was performed to identify whether the protein bound to or degraded any of the cell wall components found in either fraction [24,45]. In the presence of both chitin and chitosan, the protein was detected in the supernatant (Fig. 4a). In addition, confocal microscopy analysis with specific staining of chitin and chitosan revealed undistinguishable fluorescence signals in untreated hyphae and hyphae treated with the protein (Supplementary Figure 2). Taken together, these results led us to disregard the interaction of the protein with either of these two main cell wall components. However, in the presence of β-glucan, the protein was found in the pellet, indicating that the protein specifically binds β-glucan (Fig. 4a). This biochemical affinity of the protein for β-glucans, along with the ultrastructural changes in the outer layers of the cell wall (Fig. 2) and fragmentation of the polymers (Fig. 2c, mass spectrometry analysis), are compelling results that support the specificity of the protein for this chemical fraction of the fungal cell wall. The precipitation observed with β-glucans suggested that the protein’s action might involve structural changes or degradation of the β-glucan matrix, leading to the formation of insoluble complexes.

**Figure 4.**
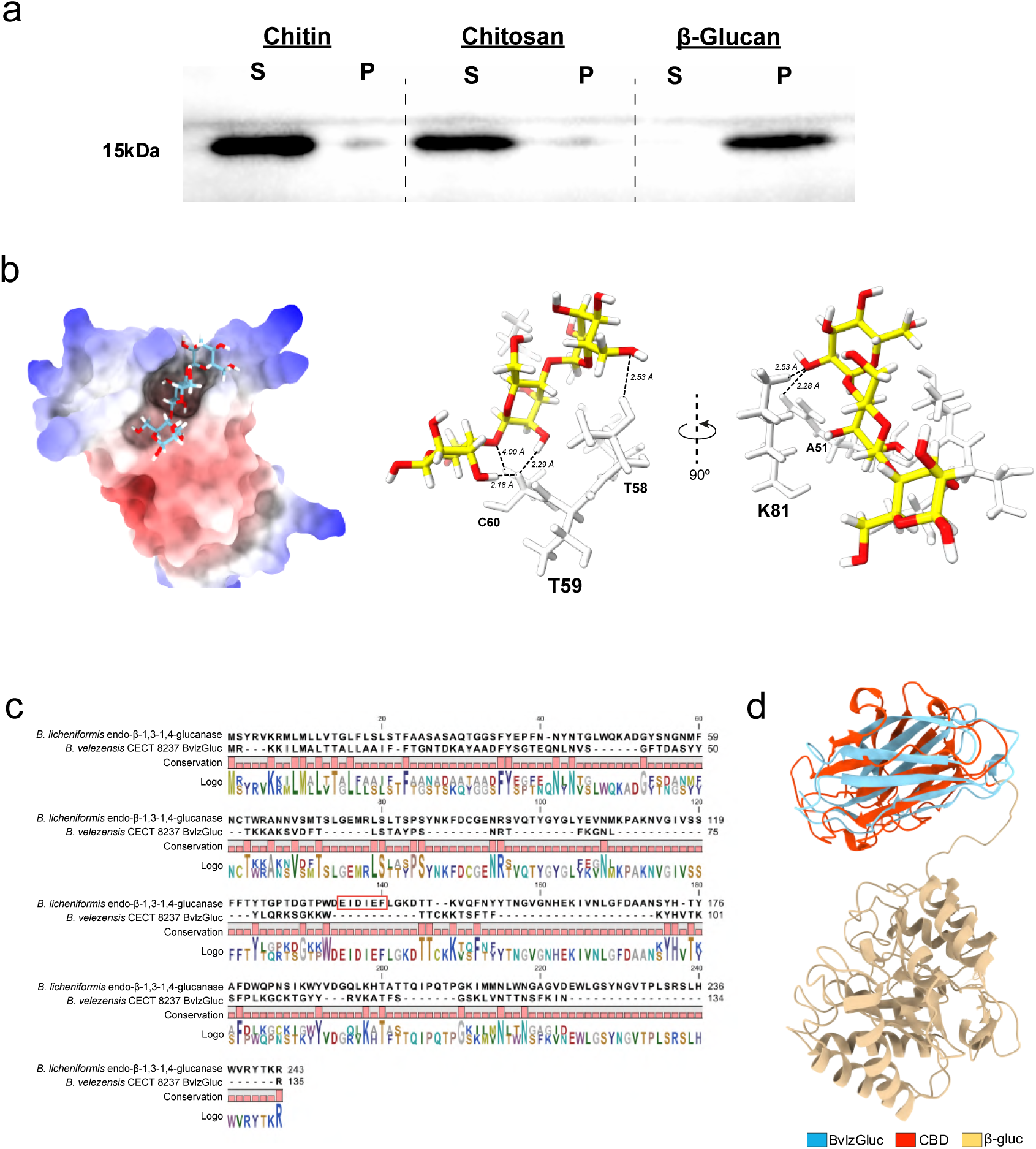
Functional analysis of BvlzGluc. **a**. Binding analysis of different carbohydrates typically present in the fungal cell wall. Through the sedimentation assay, BvlzGluc was consistently found in the pellet along with β-glucan, indicating a specific binding interaction between the enzyme and this polysaccharide. This finding suggests that BvlzGluc has a high affinity for β-glucan, likely playing a crucial role in its degradation or modification. **b**. Structural model of BvlzGluc depicted as hydrophobic potential showing the putative interaction with β-glucan. The putative binding site was formed by the cavity between β-strands β5 and β6, located at the protein core (left). Molecular docking between the β-glucan molecule and BvlzGluc. The proposed binding site is formed by residues A51, T58, T59, C60 and K81; the measured distances between β-glucan and the side chains of these residues were 2.28 Å, 2.53 Å, 2.18 Å, 2.29 Å, 4.00 Å and 2.53 Å, respectively (right). **c**. Sequence comparison between BvlzGluc from *B. velezensis* CECT 8237 and the endo-β-1,3-1,4-glucanase from *B. licheniformis* revealed significant divergence between these proteins, suggesting that BvlzGluc does not resemble canonical β-glucanases despite its ability to bind to β-glucan. Moreover, BvlzGluc lacks the conserved EIDIEF motif (red square), a hallmark of many β-glucanases, which plays a critical role in catalytic function. **d**. Comparative structural analysis of BvlzGluc and a β-glucanase from *B. velezensis* CECT 8237 revealed a striking similarity between BvlzGluc and the carbohydrate binding domain of the β-glucanase. Although the overall tertiary structures of the two proteins differ, BvlzGluc shares key structural features with the binding domain responsible for recognition and interaction with β-glucan.

Molecular docking of the protein with β-glucan was carried out to determine the potential binding site and to identify the residues involved in the interaction. The analysis predicted 31 clusters of putative binding sites distributed across 30 different regions of the protein. The most energetically favorable binding site, ΔG=-7.6 kcal mol^-1^, was located within the core of the protein. The binding of β-glucan to a protein mostly relies on the establishment of six hydrogen bonds with residues A51, T58, T59, C60, and K81. These interactions featured remarkably close distances, measuring 2.28 Å, 2.53 Å, 2.18 Å, 2.27 Å, 4.00 Å, and 2.46 Å, respectively (Fig. 4b).

On the basis of all the collected data, we classified the protein as a glycoside hydrolase. Amino acid sequence comparison with other glycoside hydrolases previously described in *B. velezensis* [46] revealed strong similarity with a protein member of glycoside hydrolase family 5 (AMQ71229), which has a structure that aligns with the defining characteristics of a canonical β-glucanase, consisting of distinct modular domains. One of these domains is a carbohydrate-binding module (CBM), which facilitates attachment to the β-glucan substrate, enhancing the affinity and specificity of the enzyme. Additionally, the structure contains a catalytic domain responsible for the hydrolysis of the β-glucan bonds, enabling the degradation of this polysaccharide [47]. The initial comparison highlighted potential functional overlaps between the two proteins; however, a more detailed analysis of their tertiary structures revealed structural dissimilarities in their catalytic domains (Fig. 4c). The similarity to the carbohydrate-binding domain points to a potential role for the protein in targeting specific polysaccharides or facilitating interactions with cell wall components, potentially assisting in the degradation process carried out by other enzymes. Additionally, the protein was compared to the endo-β-1,3-1,4-glucanase from *Bacillus licheniformis* (PDB ID: 1GBG) (Fig. 4d), the closest phylogenetically crystallized protein [48]. The analysis revealed a sequence similarity of only 12%. One of the most notable differences was the absence of the conserved amino acid motif EIDIEF, which is a hallmark feature of the *B. licheniformis* β-glucanase. This motif is often critical for the catalytic function of many β-glucanases, implying that the protein might employ a different mechanism for substrate recognition or catalysis [49,50].

### 3.6. BvlzGluc is a noncanonical β-glucanase

The phylogenetic analysis of the amino acid sequence revealed that BvlzGluc clusters with other β-glucanases from *B. subtilis* and *B. cereus* (Fig. 5a; Supplementary Table 2); despite this, it exhibited notable differences from other *B. velezensis* proteins previously described as β-glucanases [46,51]. This observation suggests that although the protein shares evolutionary ancestry with β-glucanases from related *Bacillus* species, it has undergone an evolutionary process distinct from that of the more commonly recognized β-glucanases in *B. velezensis*. The fact that BvlzGluc clusters more closely with proteins from *B. subtilis* and *B. cereus* points to a conserved evolutionary lineage between these species, highlighting potential functional similarity or shared ecological roles in these bacterial species. Furthermore, the placement of this protein within glycoside hydrolase family 16 (GH16) is of particular interest. The GH16 family is known for containing enzymes that primarily hydrolyze β-glycosidic bonds in polysaccharides, particularly β-glucans such as lichenin [52,53]. BvlzGluc features a jelly roll structure, a distinctive characteristic of the GH16 family of glucan hydrolases [48,50,54]. This architecture is composed of several antiparallel beta-sheets that contribute to its thermal stability and function, particularly through the formation of β-strands in two twisted antiparallel β-sheets, which creates a concave and convex surface (Fig. 5b). These surfaces assist in stabilizing interactions such as hydrogen bonding and hydrophobic contacts, which are crucial for maintaining structural integrity at elevated temperatures [55,56]. A prominent feature of this configuration is the presence of a deep groove on the protein’s surface, which acts as the substrate-binding site (Fig. 4b). Interestingly, this groove is absent in BvlzGluc but apparently does not affect its enzymatic functionality; therefore, BvlzGluc may utilize alternative mechanisms for substrate binding or interaction, possibly relying on different surface properties or conformational flexibility to achieve its biological role. This structural divergence from the canonical β-glucanases also suggests that this noncanonical enzyme may have evolved to fulfill a unique ecological or biological function within its environment. This could include specialized roles in the degradation of certain polysaccharides in specific niches, such as host plants, or in the processing of more complex or recalcitrant substrates that canonical β-glucanases might not efficiently process.

**Figure 5.**
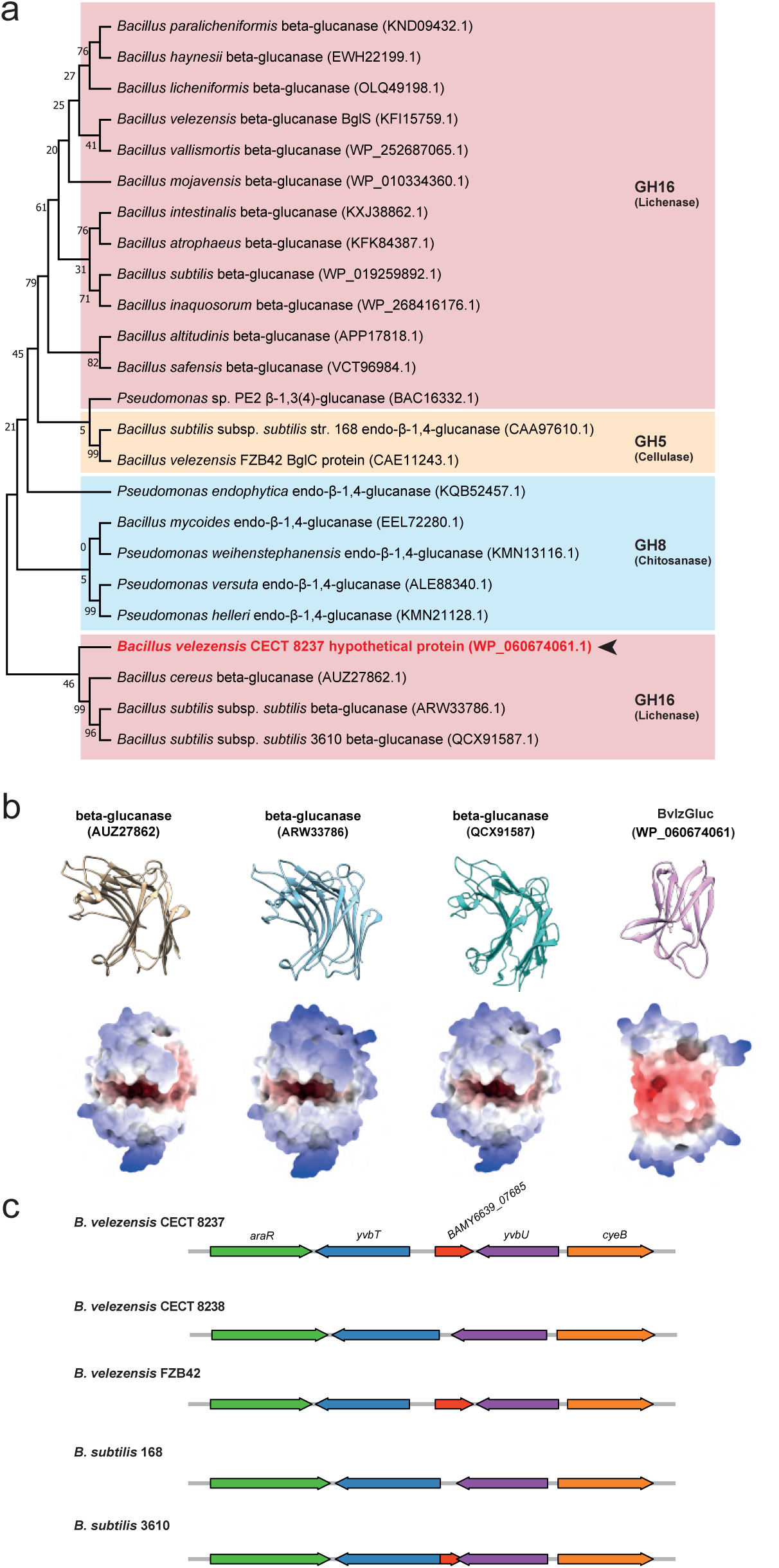
Phylogenetic and structural analysis of BvlzGluc (WP_060674061) and its structural comparison with β-glucanases from other phylogenetically related bacterial strains. **a**. In the phylogenetic analysis, BvlzGluc tended to group with β-glucanases from *B. cereus* and *B. subtilis* rather than with β-glucanases from other *B. velezensis* strains. This unexpected grouping suggests that BvlzGluc may have functional or structural characteristics more similar to those of the β-glucanases of these species. Further analysis revealed that BvlzGluc likely belongs to glycoside hydrolase family 16 (GH16), which is known for its ability to degrade complex polysaccharides containing mixed-linkage glucans, such as β-1,3- and β-1,4-glucans. Enzymes from this family specialize in degrading structural polysaccharides present in plant and fungal cell walls, highlighting the potential role of BvlzGluc in targeting a broad range of substrates. The evolutionary divergence of BvlzGluc from other *B. velezensis* β-glucanases could reflect functional adaptations, possibly providing unique substrate specificities or modes of action within the GH16 family. **b**. Structural comparison of proteins within the same phylogenetic cluster revealed that despite their evolutionary alignment, the proteins presented notable structural differences. While they all share the conserved jelly roll fold, which is a common motif in many β-barrel proteins, BvlzGluc does not possess the deep groove typically associated with enzymatic activity. This absence suggests that although these proteins may be evolutionarily related, they might have diverged in specific functions. **c**. Analysis of the genetic context of the BvlzGluc-encoding gene *BAMY6639_07685* revealed interesting patterns of distribution among *Bacillus* species. This gene was identified in *B. velezensis* FZB42, the representative strain for the group, suggesting that it plays a significant role in the functional capabilities of this strain, potentially contributing to its specialized enzymatic activities. However, in *B. subtilis*, the gene was either completely absent or truncated, indicating a possible loss or modification of function during evolution.

A comparative gene cluster analysis was conducted to determine the genetic context of the BvlzGluc protein across different *Bacillus* strains. The gene clusters for *B. velezensis* CECT 8237, CECT 8238, FZB42, and *B. subtilis* 168 and 3610 presented a high degree of structural conservation, except for the BAMY6639_07685-encoding gene. This gene is complete and functional in *B. velezensis* CECT 8237 as well as in *B. velezensis* FZB42. However, it is absent in the strains *B. velezensis* CECT 8238 and *B. subtilis* 168. Interestingly, in *B. subtilis* 3610, the gene is annotated as a pseudogene, indicating that while remnants of the gene remain, it likely does not produce a functional protein (Fig. 5c). The absence or inactivation of this gene in some strains suggests potential evolutionary divergence related to specific functional requirements or ecological adaptations. In strains in which the gene is functional, such as *B. velezensis* CECT 8237 and FZB42, the presence of BvlzGluc could be associated with distinct metabolic pathways or environmental interactions that confer a selective advantage, particularly in terms of plant‒microbe interactions or antifungal activity. In fact, comparative genomic analyses revealed that *B. velezensis* strains, including CECT 8237 and FZB42, share a core genome enriched in secondary metabolism genes, including key genes involved in plant‒microbe interactions and antifungal activities [14,57]. In contrast, the lack of this gene or the lack of nonfunctional versions reflects evolutionary divergence, suggesting an adaptive response in which the gene becomes nonessential in particular ecological niches [13,58].

### 3.7. Concentration-dependent aggregation of BvlzGluc reduces its efficacy, while its biochemical stability highlights its potential for industrial applications

BvlzGluc purified *in vitro* yielded a distinctive band of 15 kDa (Fig. 6a), which was consistent with the estimated molecular weight at a concentration of 0.5 μM. However, as the protein concentration increased to 1.5 and 2.5 μM, the 15 kDa band was replaced by a higher molecular weight band of 50 kDa. Interestingly, the decreased efficacy of the biological functionality of the protein against *Botrytis* was correlated with the increase in protein concentration (Fig. 6b). This inverse relationship suggests the formation of larger molecular weight aggregates, likely due to protein aggregation or oligomerization. To further investigate this phenomenon, the level of protein aggregation was evaluated at different concentrations. TEM analysis of negatively stained protein suspensions revealed multiple aggregates at high protein concentrations, which were absent at lower concentrations (Fig. 6c). This finding indicates that a high concentration of protein may promote intermolecular interactions, leading to the aggregation of less soluble and less functional complexes than the monomeric form does.

**Figure 6.**
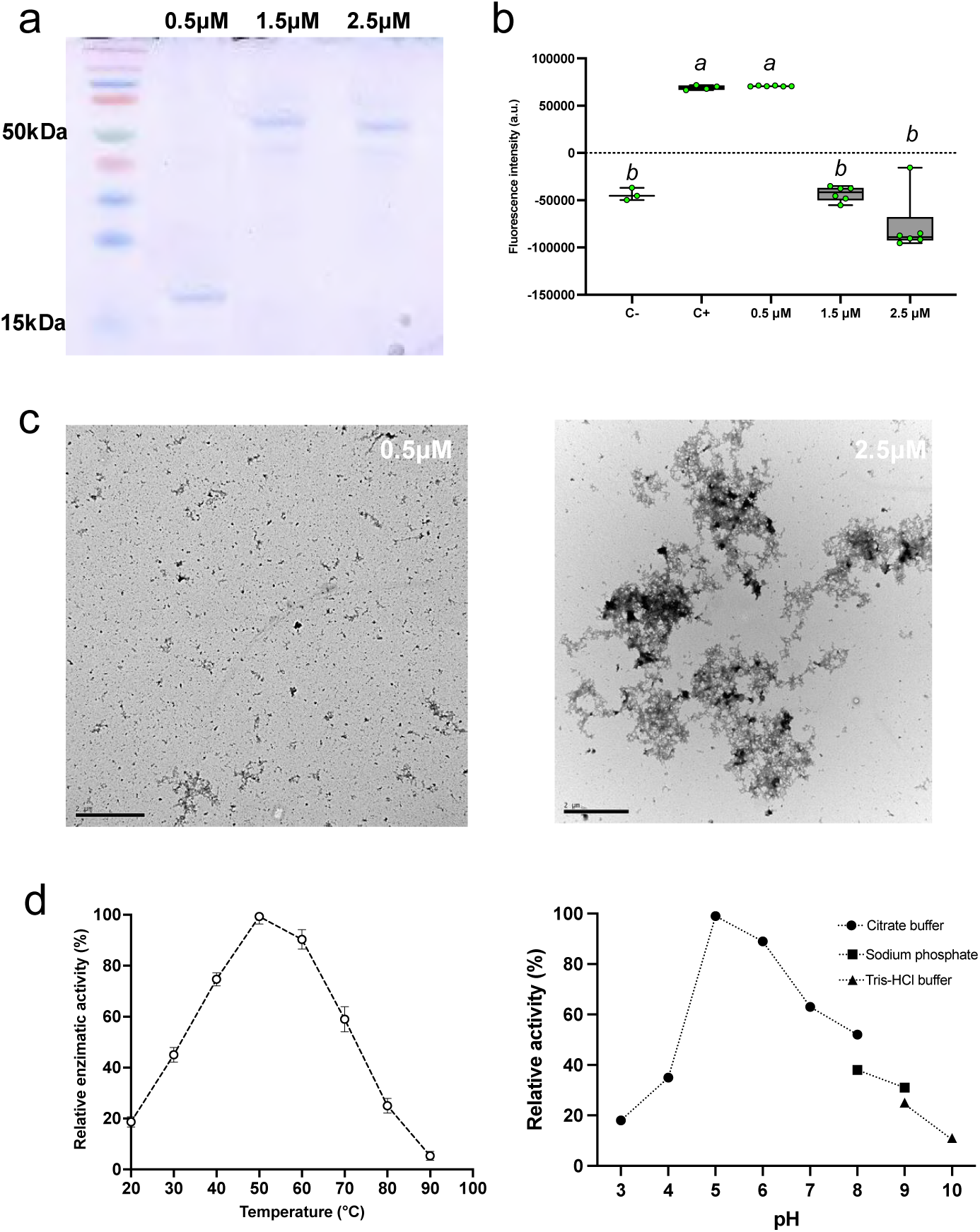
Purification and characterization of BvlzGluc. **a**. SDS‒PAGE analysis revealed an unexpected phenomenon during purification: while the protein was successfully purified (15 kDa), increasing its concentration resulted in the appearance of a band corresponding to an entity with a molecular weight higher than expected (50 kDa). This result suggests that BvlzGluc might form higher-order oligomers or complexes at elevated concentrations, which could explain the shift in molecular weight. **b**. In terms of antimicrobial activity, BvlzGluc has the ability to induce the formation of reactive oxygen species (ROS) in *Botrytis cinerea*, but this activity relies on the protein concentration. At low concentrations, the protein effectively triggered an oxidative burst, contributing to its potential role as an antimicrobial agent. However, as the concentration increased, its ability to induce ROS generation decreased, leading to a complete loss of activity at higher concentrations. **c**. Aggregation of BvlzGluc at high concentrations was confirmed, supporting the hypothesis that a lack of activity is linked to protein aggregation. This behavior could be due to the tendency of proteins to self-associate at relatively high concentrations, resulting in the formation of inactive aggregates. **d**. Physicochemical characterization of BvlzGluc revealed important details about its stability and activity under various environmental conditions. The optimal temperature for its enzymatic activity was established at 50 °C, indicating that BvlzGluc can function efficiently at relatively high temperatures, making it suitable for applications with thermal stability requirements. Additionally, the optimal pH was 5, suggesting that BvlzGluc performs best under mildly acidic conditions, which could be typical of certain biological niches or industrial processes.

Previous studies have indicated that β-glucanases can aggregate under specific conditions, resulting in larger molecular weight complexes that diminish enzymatic efficiency [59,60]. This aggregation phenomenon has been observed in various organisms, underscoring the importance of concentration and environmental factors in modulating enzymatic functionality [61,62] [60,63,64]. It has been proposed that this aggregation functions as a negative feedback mechanism to prevent the overproduction of metabolites that might be deleterious to the producer cell and thus enable the organism to maintain homeostatic balance and display broader biological processes [64,65]. The resistance of the purified protein to different physicochemical conditions, such as temperature and pH, was further tested. The optimal temperature at which the highest catalytic activity is exhibited by the enzyme was 50 °C (Fig. 6d), which highlights the outstanding thermal stability of the protein [66,67]. Additionally, the enzyme was determined to operate most effectively under mildly acidic conditions (pH 5) (Fig. 6e). This is a common feature of many β-glucanases in certain plant tissues that are crucial for plant growth and defense against pathogens, facilitating the degradation of pathogen cell walls [55,68,69]. The combination of tolerance to moderately high temperatures and a slightly acidic pH optimum indicates that the protein is likely adapted to function in specific ecological or physiological niches where these conditions prevail, such as soil or plant environments with fluctuating temperatures and pH levels [39,70]. In addition, this stability of the enzyme indicates potential for biotechnological application in biofuel production or food processing, where stable enzymatic activity at relatively high temperatures and specific pH ranges is often required [66,71,72].

## 4. Conclusion

BvlzGluc is a β-glucanase enzyme characterized by its ability to degrade the β-glucan layer of the fungal cell wall, resulting in significant structural malformations that compromise pathogen development and reduce virulence. This enzymatic degradation weakens the fungal cell wall, leading to a measurable reduction in lesion size. Overall, we conclude that BvlzGluc functions as a β-glucanase, most likely belonging to the glycoside hydrolase family 16 (GH16). However, the significant structural differences observed suggest that it represents a noncanonical member of this family. These unique structural features and enzymatic properties raise the possibility of considering BvlzGluc as part of a new subclass or even a novel family of β-glucanases. Nevertheless, further studies are necessary to confirm such a classification.

Additionally, BvlzGluc demonstrates exceptional stability under a wide range of physicochemical conditions, including temperature and pH, which further enhances its practicality for biotechnological applications. The relatively low structural complexity of BvlzGluc simplifies production processes, making it a promising candidate for scalable commercial use in agriculture and other industries. These attributes highlight the potential of this enzyme as a sustainable and cost-effective solution for pathogen control. Research into the bacterial factors involved in antagonistic interactions remains crucial to understanding the complexity of these processes and their potential applications in various fields. In this context, it is essential to delve deeper into how certain microorganisms can act antagonistically, inhibiting pathogens or enhancing the growing environment. Furthermore, the study of these mechanisms favors the efficient transfer of this knowledge into products that can be used in sustainable agriculture, promoting more eco-friendly and efficient methods.

## Acknowledgments

We thank Saray Morales Rojas for its technical support; Alicia Esteban and David Navas from the IHSM and SCAI microscopy units, respectively, for their technical support in confocal microscopy; and Mercedes Martín Rufián from the Proteomic Unit from the SCAI-UMA for their technical suggestions.

This work was partially supported by grants from the ERC Starting Grant (BacBio 637971), Plan Nacional de I+D+I of the Ministerio de Economía y Competitividad (PID2019-107724GB-I00, PID2022-141664NB-I00), research contract 8.06/60.4086 with KOPPERT B.V. (The Netherlands), and Proyecto Jóvenes Investigadores from the Plan Propio de Universidad de Málaga (B1-2021_34), with D.V.C. as the Principal Investigator. D.V.C. is funded by the Incorporación de Doctores PAIDI from Junta de Andalucía program (DOC_00266).

## Author contributions

D.R. and D.V.C. conceived the study; D.R. and D.V.C. designed the experiments; D.V.C. and A.M.C. performed the main experimental work; D.V.C. performed and designed the confocal microscopy work and data analysis; D.V.C. performed docking analyses; D.V.C. performed MALDI‒TOF analysis; D.R. and D.V.C. wrote the manuscript; and A.V. and A.P.G. contributed critically to writing the final version of the manuscript.

---

**Supplementary table 1.**
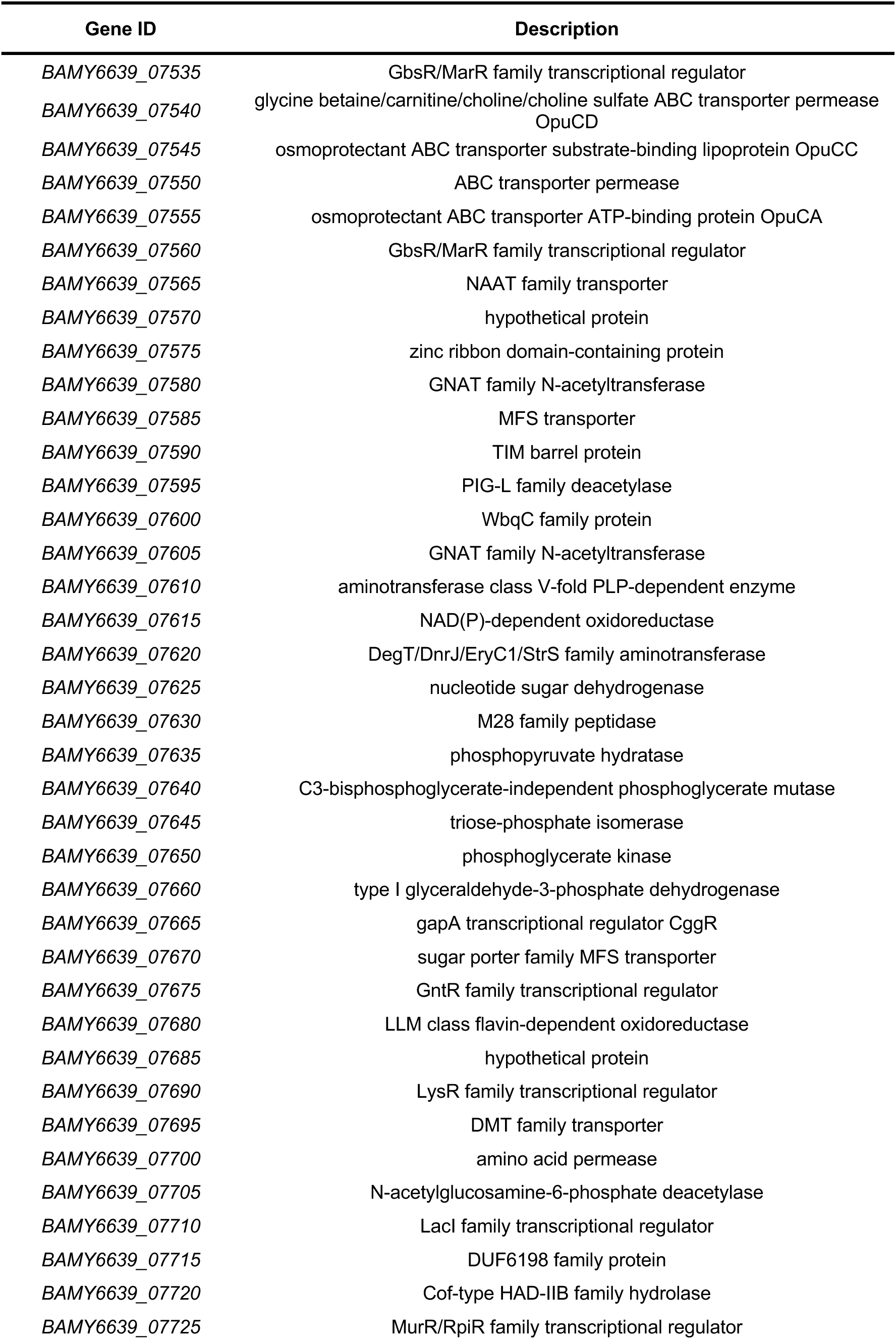
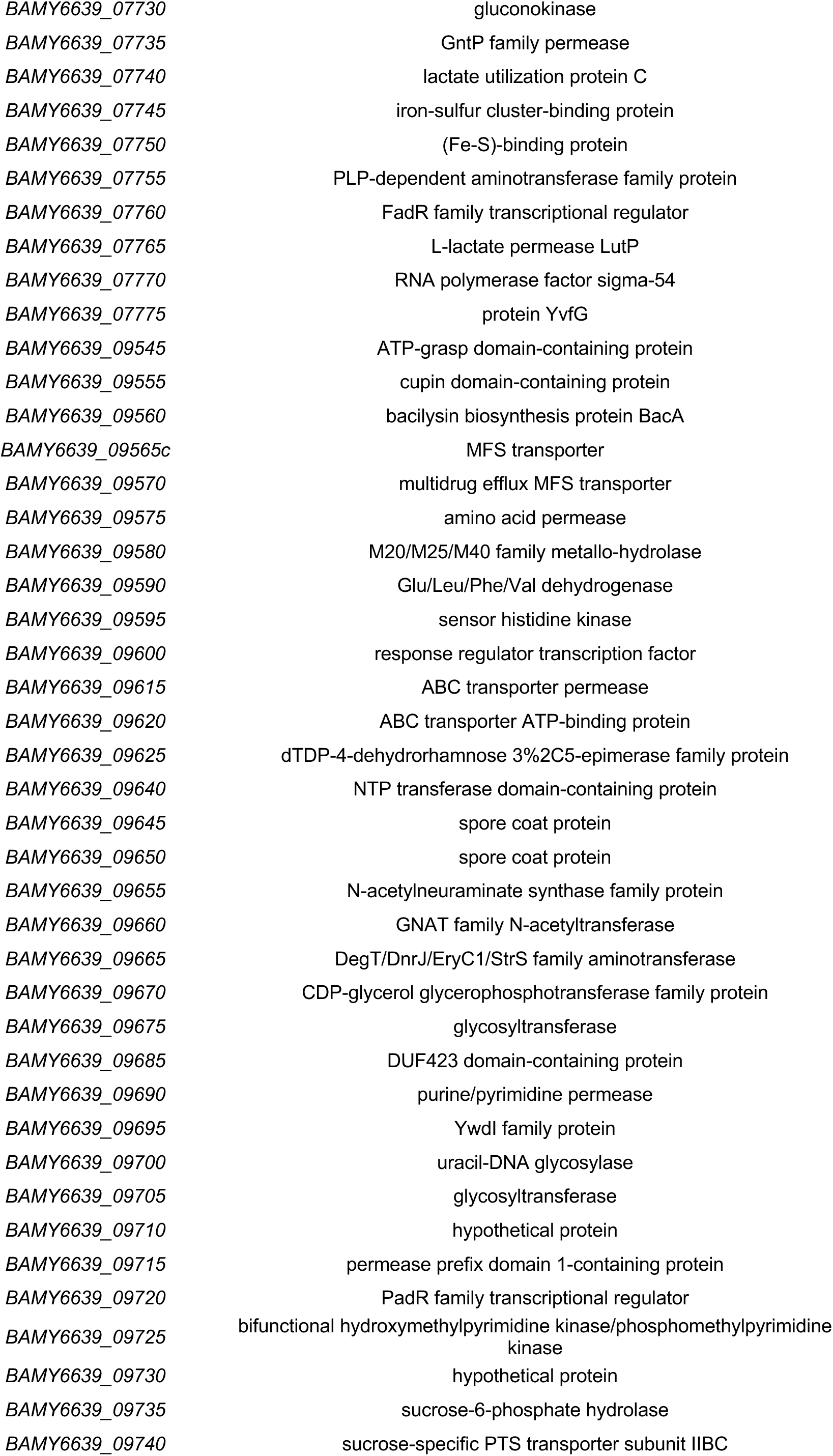
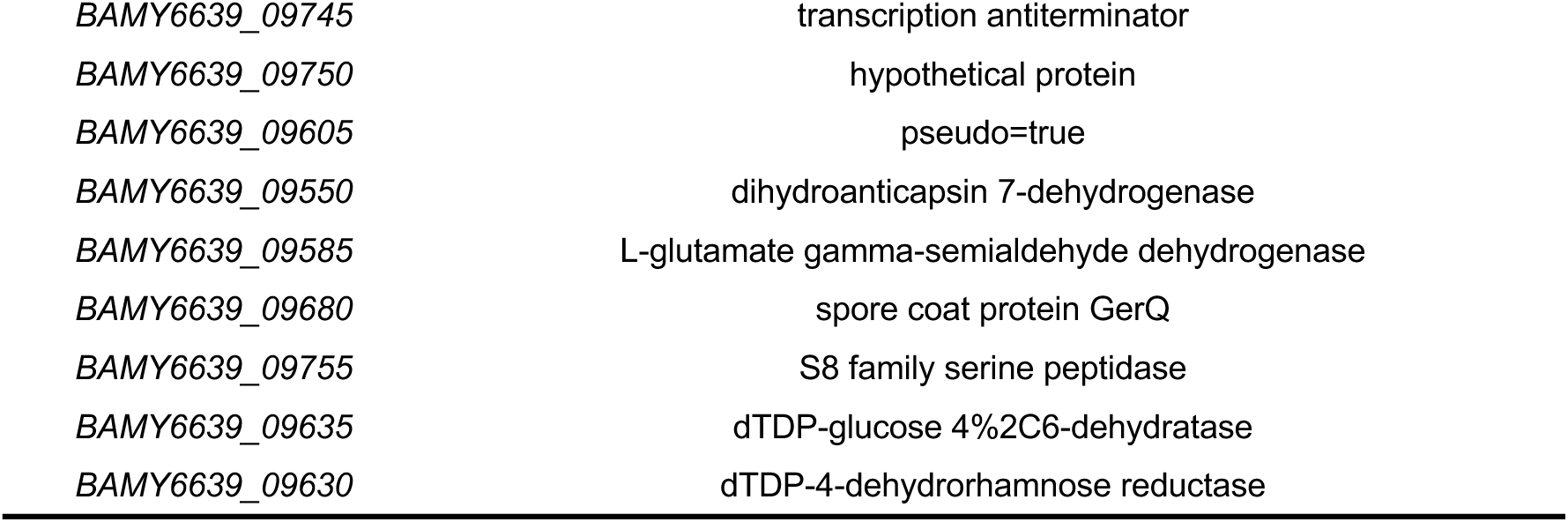
Gene functions coded in the fosmid-cloned fragments.

**Supplementary table 2.**
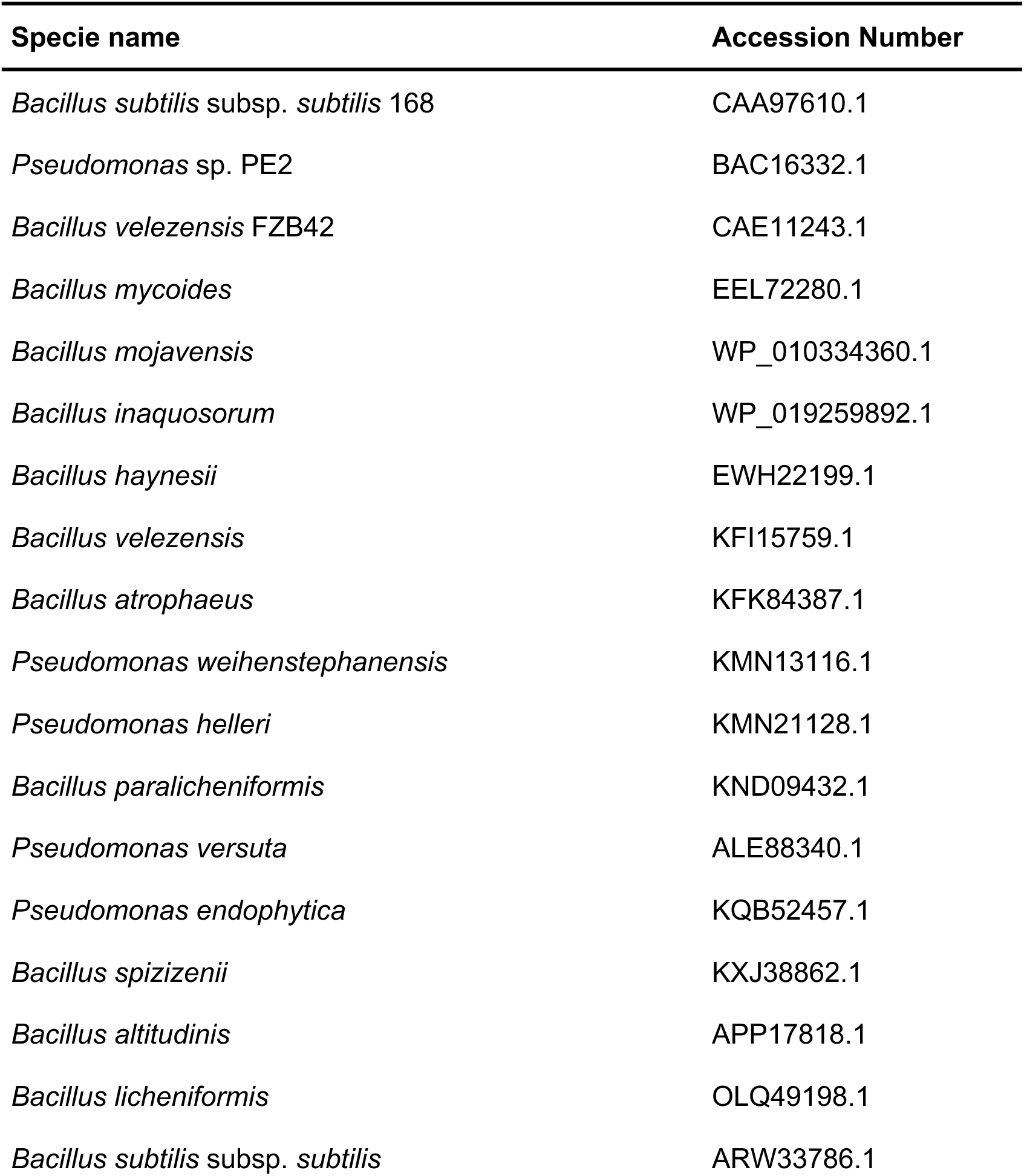
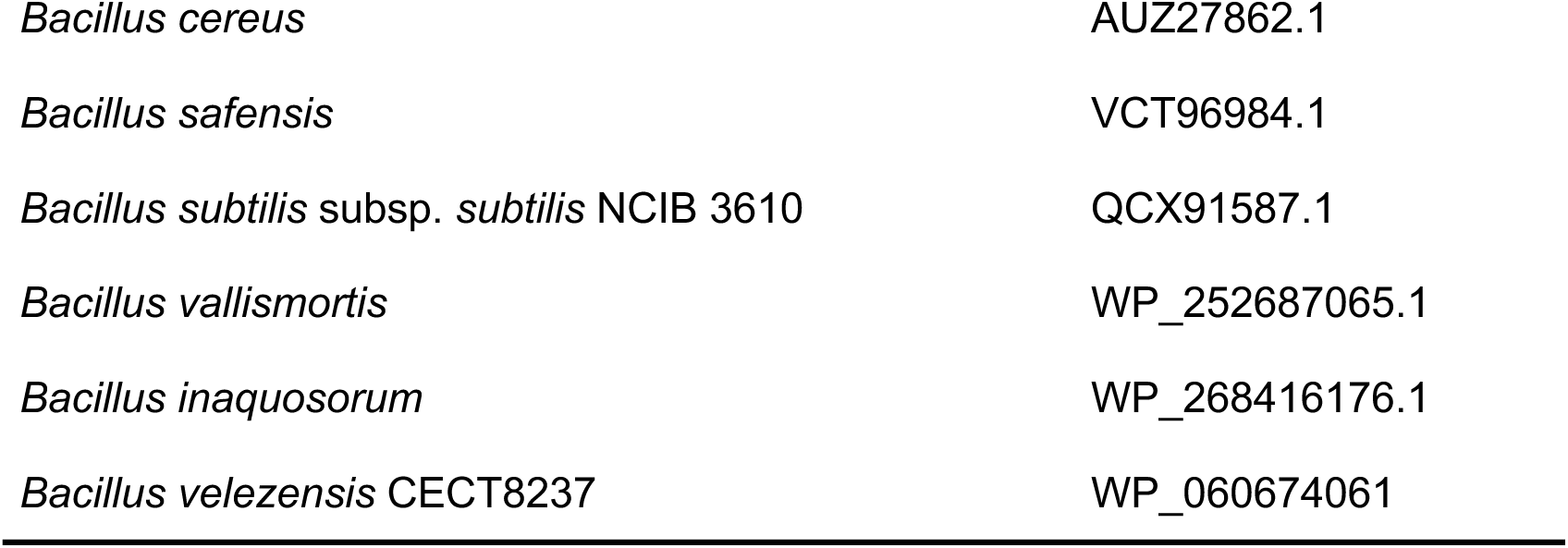
List of sequences used in this study.

## References

[1] J. Ruiz-Herrera, L. Ortiz-Castellanos, Cell wall glucans of fungi. A review, Cell Surface 5 (2019). 10.1016/j.tcsw.2019.100022.

[2] G. Camilli, G. Tabouret, J. Quintin, The Complexity of Fungal β-Glucan in Health and Disease: Effects on the Mononuclear Phagocyte System, Front Immunol 9 (2018). 10.3389/fimmu.2018.00673.

[3] J.W. Cao, Q. Deng, D.Y. Gao, B. He, S.J. Yin, L.C. Qian, J.K. Wang, Q. Wang, A novel bifunctional glucanase exhibiting high production of glucose and cellobiose from rumen bacterium, Int J Biol Macromol 173 (2021) 136–145. 10.1016/j.ijbiomac.2021.01.113.

[4] L.K. Edison, P.K. Satheeshkumar, N.S. Pradeep, Industrial Production and Purification of Recombinant Beta-Glucanases, in: N.S. Pradeep, L.K. Edison (Eds.), Microbial Beta Glucanases: Molecular Structure, Functions and Applications, Springer Nature Singapore, Singapore, 2022: pp. 171–185. 10.1007/978-981-19-6466-4_11.

[5] D. Morales, Food By-Products and Agro-Industrial Wastes as a Source of β-Glucans for the Formulation of Novel Nutraceuticals, Pharmaceuticals 16 (2023). 10.3390/ph16030460.

[6] C. Falter, C. Zwikowics, D. Eggert, A. Blümke, M. Naumann, K. Wolff, D. Ellinger, R. Reimer, C.A. Voigt, Glucanocellulosic ethanol: The undiscovered biofuel potential in energy crops and marine biomass, Sci Rep 5 (2015). 10.1038/srep13722.

[7] P.F. Ávila, M.F. Silva, M. Martins, R. Goldbeck, Cello-oligosaccharides production from lignocellulosic biomass and their emerging prebiotic applications, World J Microbiol Biotechnol 37 (2021). 10.1007/s11274-021-03041-2.

[8] J.C.G. Cortés, M.Á. Curto, V.S.D. Carvalho, P. Pérez, J.C. Ribas, The fungal cell wall as a target for the development of new antifungal therapies, Biotechnol Adv 37 (2019). 10.1016/j.biotechadv.2019.02.008.

[9] Y. Jha, Applications of Microbial Beta-Glucanase in Crop Improvement Under Biotic and Abiotic Stress, in: N.S. Pradeep, L.K. Edison (Eds.), Microbial Beta Glucanases: Molecular Structure, Functions and Applications, Springer Nature Singapore, Singapore, 2022: pp. 99–116. 10.1007/978-981-19-6466-4_7.

[10] X. Gu, Z. Cao, Z. Li, H. Yu, W. Liu, Plant immunity suppression by an β-1,3-glucanase of the maize anthracnose pathogen Colletotrichum graminicola, BMC Plant Biol 24 (2024). 10.1186/s12870-024-05053-0.

[11] T. Takashima, N. Komori, K. Uechi, T. Taira, Characterization of an antifungal β-1,3-glucanase from Ficus microcarpa latex and comparison of plant and bacterial β-1,3-glucanases for fungal cell wall β-glucan degradation, Planta 258 (2023). 10.1007/s00425-023-04271-4.

[12] J.S. Sandhu, M.K. Sidhu, I.S. Yadav, Control of Fungal Diseases in Agricultural Crops by Chitinase and Glucanase Transgenes, in: E. Lichtfouse (Ed.), Sustainable Agriculture Reviews, Springer International Publishing, Cham, 2017: pp. 163–212. 10.1007/978-3-319-48006-0_6.

[13] Q. Zeng, J. Xie, Y. Li, X. Chen, X. Gu, P. Yang, G. Hu, Q. Wang, Organization, evolution and function of fengycin biosynthesis gene clusters in the Bacillus amyloliquefaciens group, Phytopathology Research 3 (2021). 10.1186/s42483-021-00103-z.

[14] P. Xu, S. Xie, W. Liu, P. Jin, D. Wei, D.G. Yaseen, Y. Wang, W. Miao, Comparative Genomics Analysis Provides New Strategies for Bacteriostatic Ability of Bacillus velezensis HAB-2, Front Microbiol 11 (2020). 10.3389/fmicb.2020.594079.

[15] T. Xu, T. Zhu, S. Li, β-1,3-1,4-glucanase gene from bacillus velezensis zj20 exerts antifungal effect on plant pathogenic fungi, World J Microbiol Biotechnol 32 (2016). 10.1007/s11274-015-1985-0.

[16] Y. Xue, Y. Zhang, K. Huang, X. Wang, M. Xing, Q. Xu, Y. Guo, A novel biocontrol agent Bacillus velezensis K01 for management of gray mold caused by Botrytis cinerea, AMB Express 13 (2023). 10.1186/s13568-023-01596-x.

[17] V. Choub, H.B. Ajuna, S.J. Won, J.H. Moon, S.I. Choi, C.E.H. Maung, C.W. Kim, Y.S. Ahn, Antifungal activity of bacillus velezensis ce 100 against anthracnose disease (Colletotrichum gloeosporioides) and growth promotion of walnut (juglans regia L.) trees, Int J Mol Sci 22 (2021). 10.3390/ijms221910438.

[18] D. Vela-Corcía, D. Aditya Srivastava, A. Dafa-Berger, N. Rotem, O. Barda, M. Levy, MFS transporter from Botrytis cinerea provides tolerance to glucosinolate-breakdown products and is required for pathogenicity, Nat Commun 10 (2019). 10.1038/s41467-019-10860-3.

[19] D. Vela-Corcía, R. Bautista, A. De Vicente, P.D. Spanu, A. Pérez-García, De novo analysis of the epiphytic transcriptome of the cucurbit powdery mildew fungus Podosphaera xanthii and identification of candidate secreted effector proteins, PLoS One 11 (2016) 1–21. 10.1371/journal.pone.0163379.

[20] J. Jumper, R. Evans, A. Pritzel, T. Green, M. Figurnov, O. Ronneberger, K. Tunyasuvunakool, R. Bates, A. Žídek, A. Potapenko, A. Bridgland, C. Meyer, S.A.A. Kohl, A.J. Ballard, A. Cowie, B. Romera-Paredes, S. Nikolov, R. Jain, J. Adler, T. Back, S. Petersen, D. Reiman, E. Clancy, M. Zielinski, M. Steinegger, M. Pacholska, T. Berghammer, S. Bodenstein, D. Silver, O. Vinyals, A.W. Senior, K. Kavukcuoglu, P. Kohli, D. Hassabis, Highly accurate protein structure prediction with AlphaFold, Nature 596 (2021) 583–589. 10.1038/s41586-021-03819-2.

[21] A. Grosdidier, V. Zoete, O. Michielin, SwissDock, a protein-small molecule docking web service based on EADock DSS., Nucleic Acids Res 39 (2011) W270–7. 10.1093/nar/gkr366.

[22] A. Grosdidier, EADock: docking of small molecules into protein active sites with a multiobjective evolutionary optimization, Proteins: Structure, … 1025 (2007) 1010–1025. 10.1002/prot.

[23] L.G. Baker, C.A. Specht, M.J. Donlin, J.K. Lodge, Chitosan, the deacetylated form of chitin, is necessary for cell wall integrity in Cryptococcus neoformans, Eukaryot Cell 6 (2007) 855–867. 10.1128/EC.00399-06.

[24] H. Volk, K. Marton, M. Flajšman, S. Radišek, H. Tian, I. Hein, Č. Podlipnik, B.P.H.J. Thomma, K. Košmelj, B. Javornik, S. Berne, Chitin-Binding Protein of Verticillium nonalfalfae Disguises Fungus from Plant Chitinases and Suppresses Chitin-Triggered Host Immunity, Molecular Plant-Microbe Interactions 32 (2019) 1378–1390. 10.1094/MPMI-03-19-0079-R.

[25] M.J. Bailey, P. Biely, K. Poutanen, Interlaboratory testing of methods for assay of xylanase activity, 1992.

[26] R. Wang, Z. Long, X. Liang, S. Guo, N. Ning, L. Yang, X. Wang, B. Lu, J. Gao, The role of a β-1,3-1,4-glucanase derived from Bacillus amyloliquefaciens FS6 in the protection of ginseng against Botrytis cinerea and Alternaria panax, Biological Control 164 (2021). 10.1016/j.biocontrol.2021.104765.

[27] S.-I. Choi, H.-I. Lim, H.B. Ajuna, J.-H. Moon, S.-J. Won, V. Choub, J.-Y. Yun, Y. Sang Ahn, Biocontrol of fungal pathogens and growth promotion in the Korean fir (Abies koreana E.H.Wilson) seedling using Bacillus velezensis CE 100, Biological Control (2024) 105620. 10.1016/j.biocontrol.2024.105620.

[28] R.G.C. Diabankana, D.M. Afordoanyi, R.I. Safin, R.M. Nizamov, L.Z. Karimova, S.Z. Validov, Antifungal Properties, Abiotic Stress Resistance, and Biocontrol Ability of Bacillus mojavensis PS17, Curr Microbiol 78 (2021) 3124–3132. 10.1007/s00284-021-02578-7.

[29] M. Zhao, D. Liu, Z. Liang, K. Huang, X. Wu, Antagonistic activity of Bacillus subtilis CW14 and its β-glucanase against Aspergillus ochraceus, Food Control 131 (2022). 10.1016/j.foodcont.2021.108475.

[30] G. Steinberg, M.A. Peñalva, M. Riquelme, H.A. Wösten, S.D. Harris, Cell Biology of Hyphal Growth, Microbiol Spectr 5 (2017). 10.1128/microbiolspec.funk-0034-2016.

[31] J.M. Bain, M.F. Alonso, D.S. Childers, C.A. Walls, K. Mackenzie, A. Pradhan, L.E. Lewis, J. Louw, G.M. Avelar, D.E. Larcombe, M.G. Netea, N.A.R. Gow, G.D. Brown, L.P. Erwig, A.J.P. Brown, Immune cells fold and damage fungal hyphae, (n.d.). 10.1073/pnas.2020484118/-/DCSupplemental.

[32] K. Miyazawa, A. Yoshimi, A. Yoshimi, K. Abe, K. Abe, K. Abe, The mechanisms of hyphal pellet formation mediated by polysaccharides, α-1,3-glucan and galactosaminogalactan, in Aspergillus species, Fungal Biol Biotechnol 7 (2020). 10.1186/s40694-020-00101-4.

[33] A. Beauvais, C. Schmidt, S. Guadagnini, P. Roux, E. Perret, C. Henry, S. Paris, A. Mallet, M.C. Prévost, J.P. Latgé, An extracellular matrix glues together the aerial-grown hyphae of Aspergillus fumigatus, Cell Microbiol 9 (2007) 1588–1600. 10.1111/j.1462-5822.2007.00895.x.

[34] S. Liu, F. Le Mauff, D.C. Sheppard, S. Zhang, Filamentous fungal biofilms: Conserved and unique aspects of extracellular matrix composition, mechanisms of drug resistance and regulatory networks in Aspergillus fumigatus, NPJ Biofilms Microbiomes 8 (2022). 10.1038/s41522-022-00347-3.

[35] M. Lorito, C. Peterbauer, C.K. Hayes, G.E. Harman, Synergistic interaction between fungal cell wall degrading enzymes and different antifungal compounds enhances inhibition of spore germination, n.d.

[36] K.F. Mitchell, R. Zarnowski, D.R. Andes, The extracellular matrix of fungal biofilms, Adv Exp Med Biol 931 (2016) 21–35. 10.1007/5584_2016_6.

[37] D. Veličković, D. Ropartz, F. Guillon, L. Saulnier, H. Rogniaux, New insights into the structural and spatial variability of cell-wall polysaccharides during wheat grain development, as revealed through MALDI mass spectrometry imaging, J Exp Bot 65 (2014) 2079–2091. 10.1093/jxb/eru065.

[38] S.G. Grishutin, A. V. Gusakov, E.I. Dzedzyulya, A.P. Sinitsyn, A lichenase-like family 12 endo-(1→4)-β-glucanase from Aspergillus japonicus: Study of the substrate specificity and mode of action on β-glucans in comparison with other glycoside hydrolases, Carbohydr Res 341 (2006) 218–229. 10.1016/j.carres.2005.11.011.

[39] T.T. Xie, J. Shen, Z. Geng, F. Wu, Y. Dong, Z. Cui, Y. Liang, X. Ye, Antifungal characterizations of a novel endo-β-1,6-glucanase from Flavobacterium sp. NAU1659, Appl Microbiol Biotechnol 108 (2024). 10.1007/s00253-024-13269-1.

[40] T. Zhou, S. Yu, H. Xu, H. Liu, Y. Rao, Stimulating fungal cell wall integrity by exogenous β-glucanase to improve the production of fungal natural products, Appl Microbiol Biotechnol 106 (2022) 7491–7503. 10.1007/s00253-022-12224-2.

[41] D.W. Lowman, L.J. West, D.W. Bearden, M.F. Wempe, T.D. Power, H.E. Ensley, K. Haynes, D.L. Williams, M.D. Kruppa, New insights into the structure of (1→3,1→6)-β-D-glucan side chains in the Candida glabrata cell wall, PLoS One 6 (2011). 10.1371/journal.pone.0027614.

[42] C. Chaliha, M.D. Rugen, R.A. Field, E. Kalita, Glycans as modulators of plant defense against filamentous pathogens, Front Plant Sci 9 (2018). 10.3389/fpls.2018.00928.

[43] A. Sánchez-Vallet, H. Tian, L. Rodriguez-Moreno, D.J. Valkenburg, R. Saleem-Batcha, S. Wawra, A. Kombrink, L. Verhage, R. de Jonge, H.P. van Esse, A. Zuccaro, D. Croll, J.R. Mesters, B.P.H.J. Thomma, A secreted LysM effector protects fungal hyphae through chitin-dependent homodimer polymerization, PLoS Pathog 16 (2020). 10.1371/journal.ppat.1008652.

[44] N. Amarsaikhan, S.P. Templeton, Co-recognition of β-glucan and chitin and programming of adaptive immunity to Aspergillus fumigatus, Front Microbiol 6 (2015). 10.3389/fmicb.2015.00344.

[45] J. Wang, J.A. Stuckey, M.J. Wishart, J.E. Dixon, A unique carbohydrate binding domain targets the Lafora disease phosphatase to glycogen, Journal of Biological Chemistry 277 (2002) 2377–2380. 10.1074/jbc.C100686200.

[46] M.C. Magno-Perez-Bryan, P.M. Martinez-Garcia, J. Hierrezuelo, P. Rodriguez-Palenzuela, E. Arrebola, C. Ramos, A. De Vicente, A. Perez-Garcia, D. Romero, Comparative genomics within the bacillus genus reveal the singularities of two robust bacillus amyloliquefaciens biocontrol strains, Molecular Plant-Microbe Interactions 28 (2015) 1102–1116. 10.1094/MPMI-02-15-0023-R.

[47] L.K. Edison, K. Menon, N.S. Pradeep, Structure and Classification of Beta-Glucanases, in: N.S. Pradeep, L.K. Edison (Eds.), Microbial Beta Glucanases: Molecular Structure, Functions and Applications, Springer Nature Singapore, Singapore, 2022: pp. 15–32. 10.1007/978-981-19-6466-4_2.

[48] M. Hahn, J. Pons, A. Planas, E. Querol, U. Heinemann, Crystal structure of Bacillus licheniformis 1,3-1,4-β-d-glucan 4-glucanohydrolase at 1.8 Å resolution, FEBS Lett 374 (1995) 221–224. 10.1016/0014-5793(95)01111-Q.

[49] G.P. Furtado, L.F. Ribeiro, C.R. Santos, C.C. Tonoli, A.R. De Souza, R.R. Oliveira, M.T. Murakami, R.J. Ward, Biochemical and structural characterization of a β-1,3-1,4-glucanase from Bacillus subtilis 168, Process Biochemistry 46 (2011) 1202–1206. 10.1016/j.procbio.2011.01.037.

[50] G.P. Furtado, S. Carli, L.P. Meleiro, J.C.S. Salgado, R.J. Ward, Enhanced hydrolytic efficiency of an engineered CBM11-glucanase enzyme chimera against barley β-D-glucan extracts, Food Chem 365 (2021). 10.1016/j.foodchem.2021.130460.

[51] Y. Jiang, H. Xu, Y. Li, H. Liu, L. Yu, M. Qiao, G. Liu, Draft genome sequence of Bacillus subtilis strain NKYL29, an antimicrobial-peptide-producing strain from soil, Genome Announc 2 (2014). 10.1128/genomeA.01140-14.

[52] Z. Huang, G. Ni, F. Wang, X. Zhao, Y. Chen, L. Zhang, M. Qu, Characterization of a Thermostable Lichenase from Bacillus subtilis B110 and Its Effects on β-Glucan Hydrolysis, J Microbiol Biotechnol 32 (2022) 484–492. 10.4014/jmb.2111.11017.

[53] J. Wang, Y. Wang, X. Wang, D. Zhang, S. Wu, G. Zhang, Enhanced thermal stability of lichenase from Bacillus subtilis 168 by SpyTag/SpyCatcher-mediated spontaneous cyclization, Biotechnol Biofuels 9 (2016). 10.1186/s13068-016-0490-5.

[54] A.H. Teh, N.H. Fazli, G. Furusawa, Crystal structure of a neoagarobiose-producing GH16 family β-agarase from Persicobacter sp. CCB-QB2, Appl Microbiol Biotechnol 104 (2020) 633–641. 10.1007/s00253-019-10237-y.

[55] C. Caseiro, J.N.R. Dias, C.M.G. de Andrade Fontes, P. Bule, From Cancer Therapy to Winemaking: The Molecular Structure and Applications of β-Glucans and β-1, 3-Glucanases, Int J Mol Sci 23 (2022). 10.3390/ijms23063156.

[56] R.M. Yennamalli, A.J. Rader, J.D. Wolt, T.Z. Sen, Thermostability in endoglucanases is fold-specific, BMC Struct Biol 11 (2011). 10.1186/1472-6807-11-10.

[57] Q. Deng, R. Wang, D. Sun, L. Sun, Y. Wang, Y. Pu, Z. Fang, D. Xu, Y. Liu, R. Ye, S. Yin, S. Xie, R. Gooneratne, Complete Genome of Bacillus velezensis CMT-6 and Comparative Genome Analysis Reveals Lipopeptide Diversity, Biochem Genet 58 (2020) 1–15. 10.1007/s10528-019-09927-z.

[58] H. Westers, R. Dorenbos, J.M. Van Dijl, J. Kabel, T. Flanagan, K.M. Devine, F. Jude, S.J. Séror, A.C. Beekman, E. Darmon, C. Eschevins, A. De Jong, S. Bron, O.P. Kuipers, A.M. Albertini, H. Antelmann, M. Hecker, N. Zamboni, U. Sauer, C. Bruand, D.S. Ehrlich, J.C. Alonso, M. Salas, W.J. Quax, Genome Engineering Reveals Large Dispensable Regions in Bacillus subtilis, Mol Biol Evol 20 (2003) 2076–2090. 10.1093/molbev/msg219.

[59] P. Nisha, Beta-Glucanase: Diverse Bacterial Sources and its Applications, in: N.S. Pradeep, L.K. Edison (Eds.), Microbial Beta Glucanases: Molecular Structure, Functions and Applications, Springer Nature Singapore, Singapore, 2022: pp. 33–49. 10.1007/978-981-19-6466-4_3.

[60] K.R.S. Celestino, R.B. Cunha, C.R. Felix, Characterization of a β-glucanase produced by Rhizopus microsporus var. microsporus, and its potential for application in the brewing industry, BMC Biochem 7 (2006). 10.1186/1471-2091-7-23.

[61] A.C. Doxey, M.W.F. Yaish, B.A. Moffatt, M. Griffith, B.J. McConkey, Functional divergence in the Arabidopsis β-1,3-glucanase gene family inferred by phylogenetic reconstruction of expression states, Mol Biol Evol 24 (2007) 1045– 1055. 10.1093/molbev/msm024.

[62] G. Chiesa, S. Kiriakov, A.S. Khalil, Protein assembly systems in natural and synthetic biology, BMC Biol 18 (2020). 10.1186/s12915-020-0751-4.

[63] K. Martin, B.M. McDougall, S. McIlroy, Jayus, J. Chen, R.J. Seviour, Biochemistry and molecular biology of exocellular fungal β-(1,3)- and β-(1,6)-glucanases, FEMS Microbiol Rev 31 (2007) 168–192. 10.1111/j.1574-6976.2006.00055.x.

[64] J. Saarikangas, Y. Barral, Protein aggregation as a mechanism of adaptive cellular responses, Curr Genet 62 (2016) 711–724. 10.1007/s00294-016-0596-0.

[65] B.G. Poulson, K. Szczepski, J.I. Lachowicz, L. Jaremko, A.H. Emwas, M. Jaremko, Aggregation of biologically important peptides and proteins: Inhibition or acceleration depending on protein and metal ion concentrations, RSC Adv 10 (2019) 215–227. 10.1039/c9ra09350h.

[66] X. Jin, J.K. Wang, Q. Wang, Microbial β-glucanases: production, properties, and engineering, World J Microbiol Biotechnol 39 (2023). 10.1007/s11274-023-03550-2.

[67] Z. Huang, G. Ni, X. Zhao, F. Wang, M. Qu, Characterization of a GH8 β-1,4-Glucanase from Bacillus subtilis B111 and Its Saccharification Potential for Agricultural Straws, J Microbiol Biotechnol 31 (2021) 1446–1454. 10.4014/jmb.2105.05026.

[68] S. Kaur, M.K. Samota, M. Choudhary, M. Choudhary, A.K. Pandey, A. Sharma, J. Thakur, How do plants defend themselves against pathogens-Biochemical mechanisms and genetic interventions, Physiology and Molecular Biology of Plants 28 (2022) 485–504. 10.1007/s12298-022-01146-y.

[69] C. dos Santos, O.L. Franco, Pathogenesis-Related Proteins (PRs) with Enzyme Activity Activating Plant Defense Responses, Plants 12 (2023). 10.3390/plants12112226.

[70] A.K. Wani, N. Akhtar, F. Sher, A.A. Navarrete, J.H.P. Américo-Pinheiro, Microbial adaptation to different environmental conditions: molecular perspective of evolved genetic and cellular systems, Arch Microbiol 204 (2022). 10.1007/s00203-022-02757-5.

[71] S. You, T. Tu, L. Zhang, Y. Wang, H. Huang, R. Ma, P. Shi, Y. Bai, X. Su, Z. Lin, H. Luo, B. Yao, Improvement of the thermostability and catalytic efficiency of a highly active β-glucanase from Talaromyces leycettanus JCM12802 by optimizing residual charge-charge interactions, Biotechnol Biofuels 9 (2016). 10.1186/s13068-016-0544-8.

[72] L.K. Edison, N.S. Pradeep, Beta-Glucanases: An Introduction, Marketing Dynamics and Industrial Applications, in: 2022: pp. 1–14. 10.1007/978-981-19-6466-4_1.

