## Supplementary material for "Characterization of BvlzGluc, a Novel Antifungal β-Glucanase from *Bacillus velezensis*, with Potential Agricultural and Industrial Applications": Supplmentary figures 1 and 2

### Slide 1
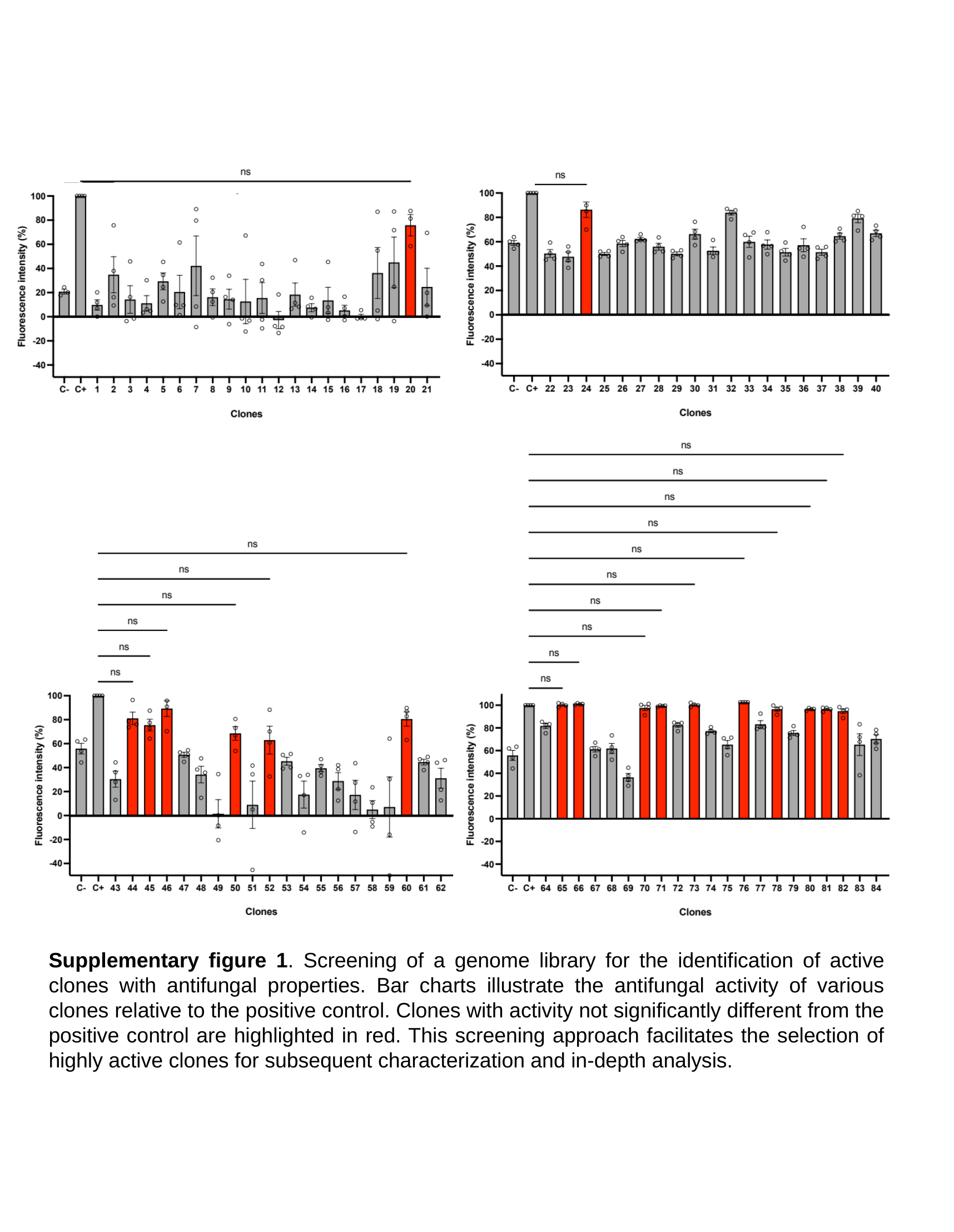

Supplementary figure 1. Screening of a genome library for the identification of active clones with antifungal properties. Bar charts illustrate the antifungal activity of various clones relative to the positive control. Clones with activity not significantly different from the positive control are highlighted in red. This screening approach facilitates the selection of highly active clones for subsequent characterization and in-depth analysis.

### Slide 2
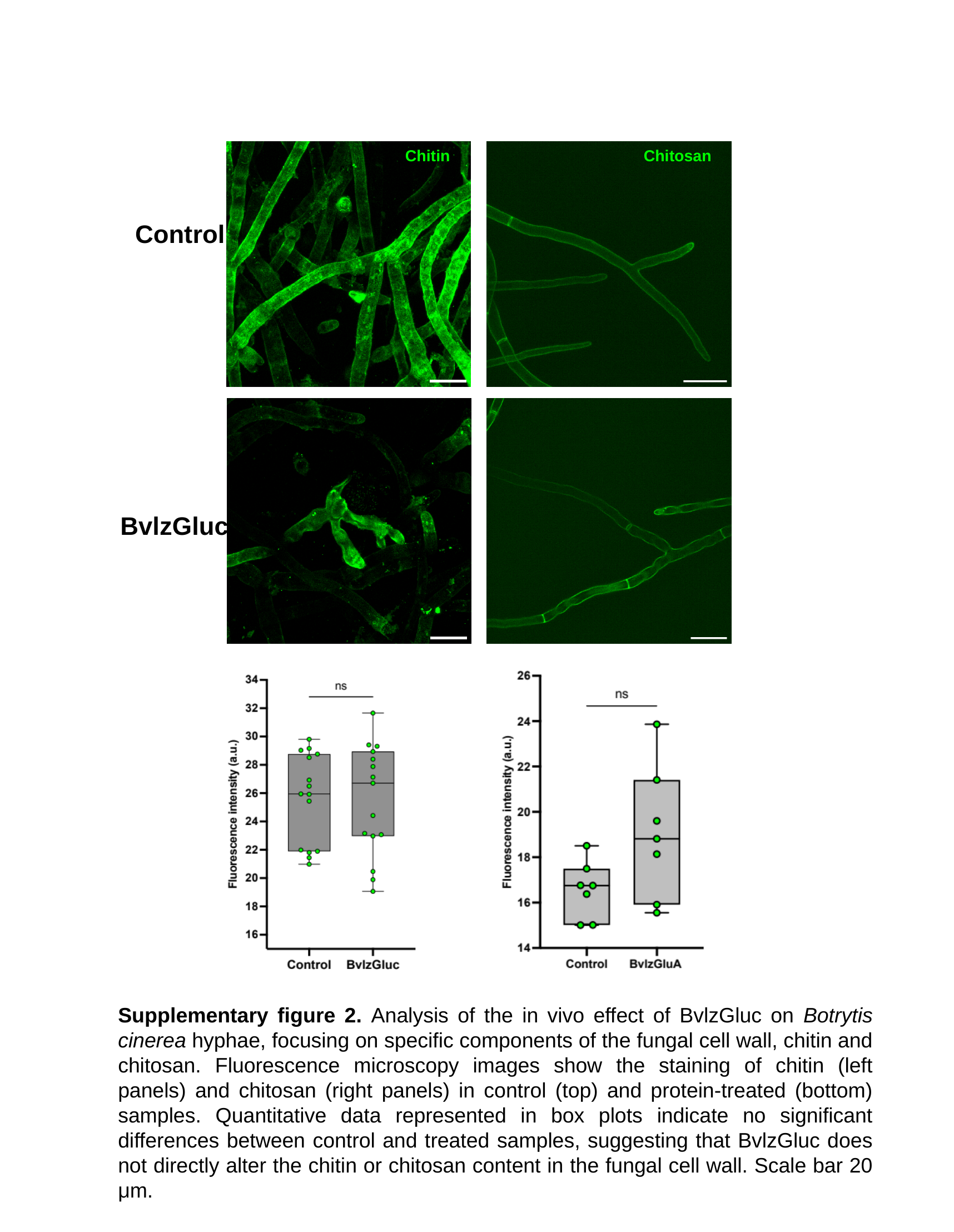

Chitin
Chitosan
Control
BvlzGluc
Supplementary figure 2. Analysis of the in vivo effect of BvlzGluc on Botrytis cinerea hyphae, focusing on specific components of the fungal cell wall, chitin and chitosan. Fluorescence microscopy images show the staining of chitin (left panels) and chitosan (right panels) in control (top) and protein-treated (bottom) samples. Quantitative data represented in box plots indicate no significant differences between control and treated samples, suggesting that BvlzGluc does not directly alter the chitin or chitosan content in the fungal cell wall. Scale bar 20 μm.
